# Effects of central dogma processes on the compaction and segregation of bacterial nucleoids

**DOI:** 10.1101/2025.07.21.665959

**Authors:** Mu-Hung Chang, Maxim O. Lavrentovich, Jan-Michael Y. Carrillo, Jaan Männik

## Abstract

The bacterial cytoplasm is characterized by a distinctive membrane-less organelle, the nucleoid, which harbors chromosomal DNA. We investigate the effects of dynamic processes associated with transcription and translation on the structure of this organelle, using coarsegrained molecular dynamics (MD) simulations implemented with out-of-equilibrium reactions. Our model captures the scale of the entire cell and incorporates a reaction-diffusion system for ribosomes and polyribosomes, combining their out-of-equilibrium dynamics with excluded volume interactions with DNA. Our findings demonstrate that out-of-equilibrium reactions increase the size of the nucleoid. In addition, we show that the nucleoid size increase is proportional to transcriptional activity. Our model reproduces the time-dependent change in nucleoid size observed in rifampicin treatment experiments, where the pool of polyribosomes is depleted. Furthermore, we find these active processes are essential for complete sister chromosome separation and correct nucleoid positioning within the cell. Overall, our study reveals the effects of the central dogma processes on the internal organization and localization of bacterial nucleoids.

**Significance:** Understanding how bacteria organize their chromosomes is fundamental to cell biology. Through our coarse-grained molecular dynamics simulations incorporating out-of-equilibrium processes of transcription and translation, we are able to capture the effects of these central dogma processes on DNA organization and demonstrate that these active biological processes expand the nucleoid and facilitate the separation of daughter chromosomes. Our simulations are compared to experimental measurements and highlight the impact of the out-of-equilibrium conditions of the living cell. These findings point out the crucial interplay between physical forces and biological activity in cellular organization, suggesting that cellular structure depends on non-equilibrium processes.

## Introduction

Unlike eukaryotic cells, bacterial cells lack a membrane-enclosed nucleus (1). Despite the lack of a nuclear membrane, bacterial chromosomal DNA forms a nucleus-like structure called the nucleoid. The nucleoid can be considered a distinct physical phase within the cell (2, 3). In the *Escherichia coli* (*E. coli*) model organism, this phase occupies about 50% of the cytosolic volume (4), with the precise fraction varying with growth conditions and bacterial species (5). Different molecules and processes have been associated with nucleoid size control, including passive DNA binding proteins (6), active SMC (structural maintenance complex) proteins (7, 8), supercoiling (9), transertion linkages that connect chromosomal DNA to the cell membrane (10), and transcription (11). However, both *in vitro* experiments with bacterial DNA (12) and *in vivo* measurements of log-phase cells (4, 13) suggest that the dominant effect on the compaction of chromosomal DNA into a nucleoid comes from macromolecular crowding. The crowder-induced nucleoid compaction has been explained using an equilibrium free energy approach based on polymer physics (3, 14, 15). In this free energy approach, DNA and crowders interact via repulsive (steric) interactions. The free energy approach predicts a segregative phase-separation of chromosomal DNA to a distinct nucleoid phase at some critical volume fraction of crowders. This phase-separated nucleoid is depleted of crowders, while the remainder of the cytosol, which we will refer to as the cytosolic phase, is enriched by them. Extensive computational simulations, where DNA is modeled as a bead on a string polymer and crowders as repulsive spheres, have confirmed these predictions (16, 17, 18, 19, 20, 21).

Soluble cytosolic proteins and ribosomes are the main macromolecular crowders in a bacterial cell that affect nucleoid compaction (4, 15). Previous equilibrium-based modeling using a free energy approach predicts that soluble proteins and ribosomes have approximately equal effects on nucleoid compaction (15). From the point of view of nucleoid compaction, proteins in the cytosol act as passive crowders that interact with DNA via excluded volume interactions. On the other hand, the majority of ribosomes in log-phase *E. coli* (*>* 80%) are assembled into polyribosomes (polysomes) (22) and the polysome dynamics will impact the nucleoid compaction. A polysome is a polymer chain of ribosomes assembled on a single mRNA molecule. This assembly already forms at the time of mRNA synthesis (transcription) and gives rise to co-transcriptional translation of proteins (23, 24). mRNA synthesis leading to the formation of polysomes is an inherently out-of-equilibrium process consuming energy. The assembly of a 70S ribosome particle on mRNA also consumes energy, requiring two molecules of GTP per ribosome assembly. Two molecules of GTP are also consumed with each step of translation, until the 70S unbinds from the mRNA and dissociates into 30S and 50S subunits. Polysomes interact strongly with chromosomal DNA during co-transcriptional translation, when they are attached to DNA via RNA polymerase (RNAP). One can thus expect that the non-equilibrium effects arising from the dynamics of polysomes significantly affect the compaction and organization of the nucleoid.

Computational studies of crowding effects on nucleoid compaction have largely been limited, so far, to equilibrium considerations. In the first part of our study, we explore the effects of non-equilibrium ribosome dynamics on the compaction and organization of the nucleoid via coarse-grained molecular dynamics simulations. We develop an out-of-equilibrium model in LAMMPS (25) making use of its *REACTER* module (26) to simulate co-transcriptional polysome assembly, translation, and polysome disassembly upon degradation of mRNA. The model also accounts for the excluded volume interactions between DNA segments, polyribosomes, and free ribosome particles. Our simulations reveal that the central dogma processes increase the nucleoid size: The dimensions of the nucleoid increase proportionally to the total transcriptional activity in the cell. The model matches experimental time-dependent data from rifampicin (Rif)-treated cells, where transcriptional activity is abruptly halted.

In the second part of this study, we extend the model to two chromosomes to address how daughter nucleoids segregate from each other. Unlike eukaryotic cells, bacteria lack a mitotic spindle and the DNA segregation mechanism is not completely understood. It has been proposed that two long DNA polymers in cylindrical confinement segregate from each other due to a force that arises from the configurational entropy of these polymers (27, 28). However, this so-called entropic mechanism cannot explain the segregation process in the early stages of DNA replication (29, 30) as the mechanism is only effective when about 50 % of the DNA is already replicated (29). Furthermore, the entropic force vanishes upon loss of physical contact between the two chromosomes (31). Therefore the entropic force also fails to explain the segregation in late cell cycle stages and in filamentous cells where the daughter chromosomes lose physical contact.

The experiments with filamentous cells show that two daughter nucleoids separate from each other over large distances (*>* 10 *µ*m) and localize robustly to the cell’s quarter positions (32). To explain this localization pattern, Miangalorra *et al.* developed a continuum reaction-diffusion model that includes central dogma reactions involving ribosomes and mRNA, and explored their effect on the separation of two nucleoids (33). The model, which included steric interactions among these species and DNA, was able to explain the experimentally observed positioning of the nucleoids in filamentous cells. Similar reaction-diffusion models were also developed and tested relative to experimental data on normal cells (34, 35). These comparisons show that an active mRNA and ribosome dynamics play a role in separating the daughter chromosomes in normal cells too, although it appears not to be the sole mechanism the cells employ.

The models by Miangalorra *et al.* (33), and in related works (34, 35) treat DNA, polysomes, and free ribosome particle densities as continuum fields. Although treating polysomes and ribosome particles as continuum fields is justifiable due to their relatively large number, it is less clear whether the approximation is justified for the DNA, which is a single connected molecule. Furthermore, all the above-mentioned models are developed in one spatial dimension, with density variations occurring only along the long axis of the cell. However, ribosomal and DNA densities along the radial direction of *E. coli* cells are known to vary from experiments (36, 37). In a 3D geometry, unlike in 1D, there is a possibility that ribosomes that preferentially accumulate at the mid-cell due to the reactions are able to diffuse past the nucleoids along the layer near the cell envelope where the DNA density is low (33). In the second part of this study, we implement simplified central dogma reactions in a 3D setting for a cell containing two chromosomes with explicit chain connectivity. We find that coupled ribosome dynamics have a significant effect on nucleoid separation in our 3D simulation, too, thus validating the main conclusions from the earlier 1D models (33, 34, 35). However, our work points out that there are significant fluctuations of DNA and ribosomes at the midcell during the initial stages of segregation - an aspect the 1D continuum models fail to predict.

## Methods

We simulate an *E. coli* cell containing one or two circular DNA molecules, along with polysomes, free ribosomal particles, and RNA polymerases (RNAPs). We do not separately model 30S and 50S ribosomal subunits for simplicity but only consider one size of free ribosomal particles that correspond to fully assembled ribosomes (70S particles). We also do not explicitly model mRNA, but implement its effects via the assembly and disassembly of polysomes. All these species interact with each other via excluded volume interactions in a cell that has a cylindrical shape with hemispherical end caps (Fig. 1a). Free ribosomal particles, polysomes, RNAPs, and DNA undergo binding and unbinding reactions. RNAP binding to DNA initiates the assembly of a new polysome. When a polysome attached to DNA via RNAP reaches a specific size (as given by the average mRNA template size), it detaches from the DNA and becomes a “free” polysome. Ribosomes attach to free polysomes from one end and detach from the other end as the translation completes. Additionally, we implement a reaction where free polysomes dissociate into ribosomal particles, which captures the effects of the degradation of the mRNA template that holds ribosomal particles in a polysome together. All reactions considered here conserve the total number of ribosomal particles (i.e., the sum of free ribosomes and polysome-bound ribosomes, *n*_ribo_, remains constant). Our simulations represent *E. coli* cells in slow-growth conditions. We conduct three simulations with different random number seeds for each setting unless otherwise specified.

**Figure 1:**
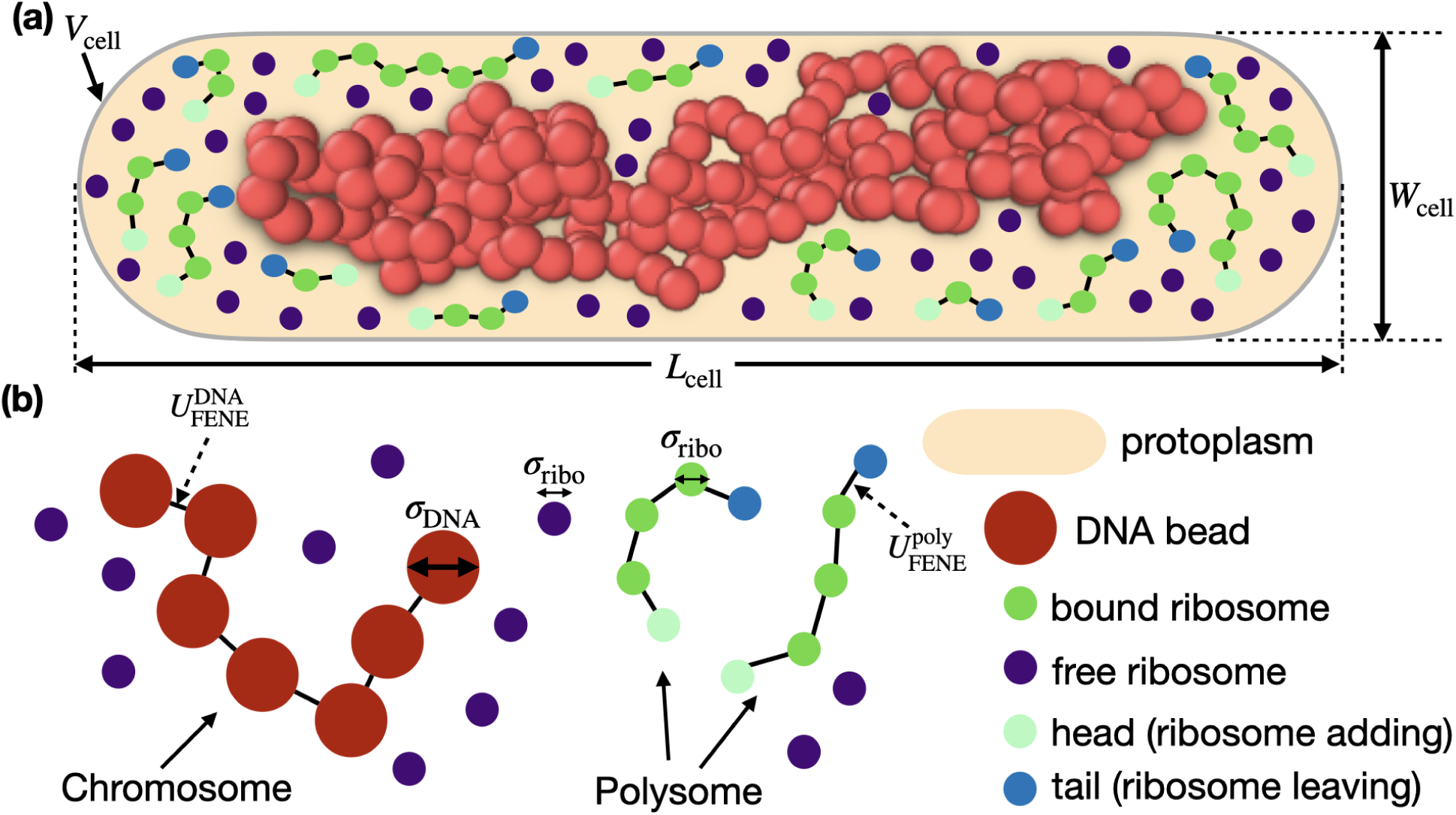
Model setup: components and potenitals. (a) A schematic of the simulated cell. The circular DNA molecule (red, shown as a snapshot from simulation) surrounded by polysomes and free ribosome particles (purple) is placed in cylindrical cell with hemispherical end caps representing the cytoplasm of an*E. coli* cell. The length and width of the domain are denoted by *L*_cell_ and *W*_cell_, respectively. *W*_cell_ also represents the diameter of the cylindrical region. For clarity, RNAPs are not shown in this schematic. (b) Key components and interactions of the coarse-grained model: DNA monomers (red) are connected by FENE bonds with potential energy 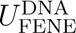 (see Eq. 1) to form the chromosome. Polysomes consist of bound ribosomes particles (green) connected by FENE bonds with potential energy 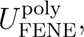 and distinct head (light green, site of ribosome addition) and tail (blue, site of ribosome removal) ribosome particles. The diameters of DNA monomers and ribosome particles are denoted as *σ*_DNA_ and *σ*_ribo_, respectively (See Methods and Fig. 2 for the details of out-equilibrium processes).

### Interaction potentials

We treat the chromosomal DNA as a ring polymer consisting of “beads on a string” (Fig. 1b). A single circular chromosome in our model is represented by 200 monomers. A similar representation of the *E. coli* chromosome has been used in previous works (4, 16). Each monomer (bead) represents a coarse-grained structural unit of the bacterial chromosome consisting of about 20 kb. The potential between adjacent beads (the “string”) is a finite extensible nonlinear elastic (FENE) potential, given by

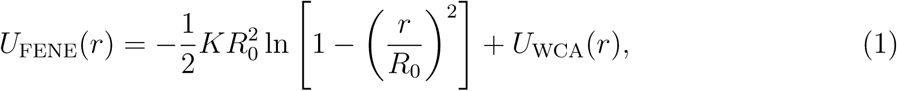

where *r* is the center-to-center distance between two adjacent beads, *K* is the effective spring constant, and *R*_0_ is the maximum distance between two beads. The second term in Eq. 1, *U*_WCA_(*r*), describes the short-ranged repulsive interaction between polymer beads. This interaction is given by the Weeks-Chandler-Andersen (WCA) potential (a truncated and shifted Lennard-Jones potential), which reads

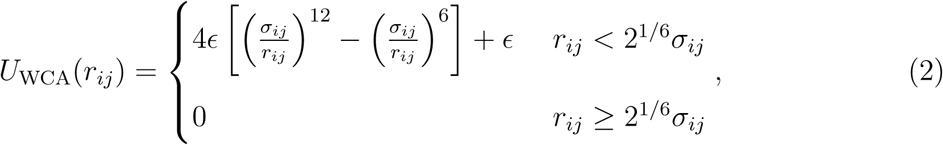

where *r_ij_* is the center-to-center distance between particles *i* and *j* and *ɛ* is a potential strength parameter. The WCA interaction is also used for the other non-bonded interactions between DNA and ribosomes and between two ribosomes. Note that the non-bonded ribosome interactions do not depend on whether or not the ribosomes are incorporated into polysomes. We choose the repulsion distance *σ_ij_* in the WCA potential via *σ_ij_* = (1*/*2)(*σ_ii_* + *σ_jj_*) where 2^1^*^/^*^6^*σ_ii_* is the effective diameter of particle *i*. However, for DNA and polysome interactions, we increase the repulsion distance by setting *σ_ij_* = 3.5*σ*_ribo_ *>* (*σ*_ribo_ + *σ*_DNA_)*/*2. This choice is informed by the previous free-energy-based study of nucleoid compaction in *E. coli* cells (15) which posits that the large size of the polysome prevents it from entering the supercoiled segments of the DNA. The resulting excluded volume between DNA and ribosomes that make up a polysome is thus expected to be larger than the excluded volume between DNA and disjoint ribosome particles.

An RNAP is connected to the DNA via a FENE potential. During co-transcriptional translation, a ribosome attaches to the RNAP via a FENE potential, as well. This potential also connects individual ribosome particles in polysomes (Fig. 1b). Unlike for the other particles, we do not include the excluded volume interactions between RNAPs and other particles because we consider RNAP to be part of the DNA monomer, which already has a large size.

All particles interact with the cell walls via the 9-3 Lennard-Jones potential (38):

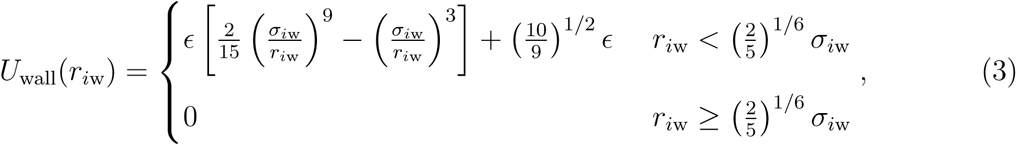

where *r_i_*_w_ is the distance between the particle and the wall surface and *σ_i_*_w_ is a length parameter, which we set to 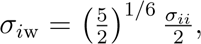 ensuring that particle *i* experiences a repulsive force when the distance between the wall and the particle is less than the radius of the particle (i.e., *r_i_*_w_ *< σ_ii_/*2). The parameters of particle-particle and particle-wall interactions used in this work are summarized in Table 1.

**Table 1:**
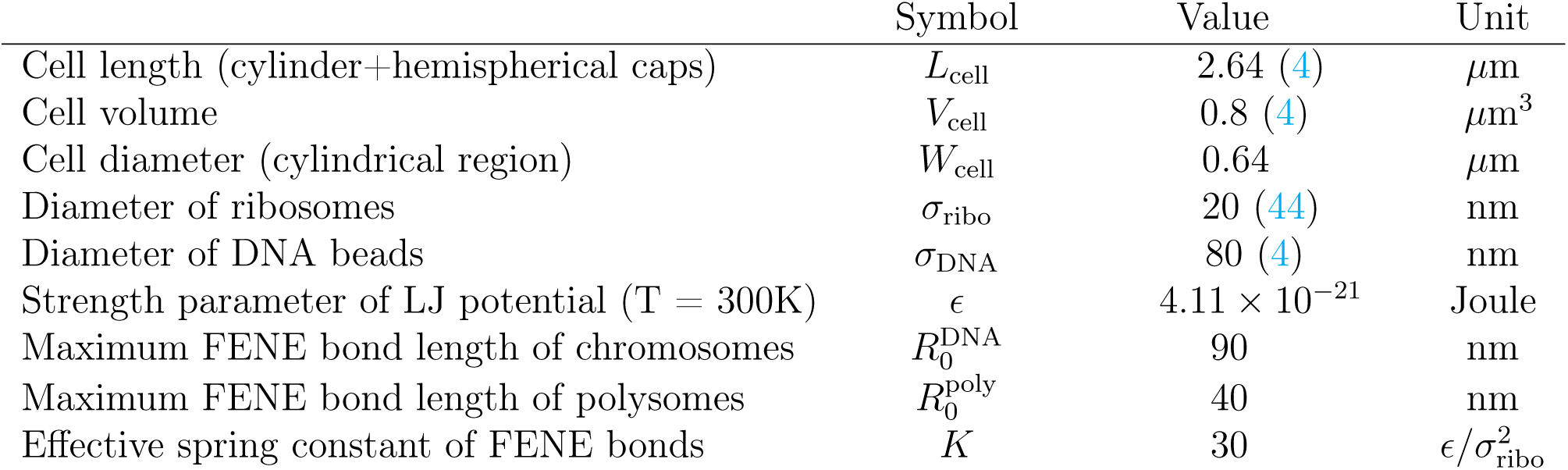
Parameters for simulation boundary, particle-particle and particle-wall interactions.

|  | Symbol | Value | Unit |
| --- | --- | --- | --- |
| Cell length (cylinder+hemispherical caps) | $L_{\text{cell}}$ | 2.64 (4) | $\mu\text{m}$ |
| Cell volume | $V_{\text{cell}}$ | 0.8 (4) | $\mu\text{m}^3$ |
| Cell diameter (cylindrical region) | $W_{\text{cell}}$ | 0.64 | $\mu\text{m}$ |
| Diameter of ribosomes | $\sigma_{\text{ribo}}$ | 20 (44) | nm |
| Diameter of DNA beads | $\sigma_{\text{DNA}}$ | 80 (4) | nm |
| Strength parameter of LJ potential ($T = 300\text{K}$) | $\epsilon$ | $4.11 \times 10^{-21}$ | Joule |
| Maximum FENE bond length of chromosomes | $R_0^{\text{DNA}}$ | 90 | nm |
| Maximum FENE bond length of polysomes | $R_0^{\text{poly}}$ | 40 | nm |
| Effective spring constant of FENE bonds | $K$ | 30 | $\epsilon/\sigma_{\text{ribo}}^2$ |

### Implementation of reactions

We employ the LAMMPS module *REACTER* (26) to implement central-dogma-related processes. *REACTER* specifies the occurrence of a reaction event *i* of two “initiator” particles *l* and *k* by three parameters: the maximum reaction distance 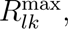 the acceptance probability of the reaction *P_i_*, and the number of LAMMPS timesteps between each attempted reaction, *N_i,_*_every_. The reaction can thus occur in every *N_i,_*_every_ time-step interval with probability *P_i_* if the positions of the two initiators, *l* and *k*, are closer to each than 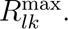 We set the reaction distance to the maximum interaction range of the repulsive interactions between particles 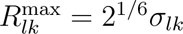 (see the WCA potential in Eq. 2). We relate the probability *P_i_* by an effective reaction rate *k_i_*, via

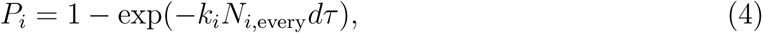

where *dτ* is the size of the LAMMPS timestep. We treat the unimolecular reactions in our model similarly, with the only difference being that the maximum reaction distance does not affect the occurrence of these reactions. For a given reaction, we determine the rates *k_i_* using their values reported in literature or estimate them by physical considerations, as will be explained in the next section. Then we choose *P_i_* according to the relation Eq. 4 in such a way that *P_i_* (and consequently, *N_i,_*_every_) is small enough to avoid synchronization of reaction events, but large enough to avoid frequently testing for the reaction condition arising from the small value of *N_i,_*_every_, which is computationally expensive and decreases the simulation efficiency. Table 2 lists chosen parameter values for different reactions.

**Table 2:** Parameters for out-of-equilibrium processes.

|  | Symbol | Value | Unit |
| --- | --- | --- | --- |
| polysome (mRNA) degradation rate | $k_{\text{deg}}$ | $6.7 \times 10^{-3}$ | 1/s |
| Ribosome unbinding rate | $k_{\text{off}}$ | 0.14 | 1/s |
| Ribosome binding rate | $k_{\text{on}}$ | $9.3 \times 10^{-5}$ | $\mu\text{m}^3/\text{s}$ |
| DNA-Bound polysomes detachment rate | $k_{\text{detach}}$ | $9.3 \times 10^{-5}$ | $\mu\text{m}^3/\text{s}$ |
| Acceptance probability of polysome (mRNA) degradation | $P_{\text{deg}}$ | 0.005 | |
| Acceptance probability of ribosome unbinding | $P_{\text{off}}$ | 0.003 | |
| Acceptance probability of ribosome binding | $P_{\text{on}}$ | 0.02 | |
| Acceptance probability of DNA-bound polysomes detachment | $P_{\text{detach}}$ | 0.02 | |
| Timestep interval of attempting polysome (mRNA) degradation | $N_{\text{deg, every}}$ | $1.5 \times 10^6$ | |
| Timestep interval of attempting ribosome unbinding | $N_{\text{off, every}}$ | $5 \times 10^4$ | |
| Timestep interval of attempting ribosome binding | $N_{\text{on, every}}$ | $2 \times 10^4$ | |
| Timestep interval of attempting DNA-bound polysome detachment | $N_{\text{detach, every}}$ | $2 \times 10^4$ | |

#### Formation of polysomes

Polysomes form in our simulations during co-transcriptional assembly. While *E. coli* contains approximately 1500-5000 RNAPs per cell (39, 40), single-molecule tracking studies reveal that about half of these are actively engaged in transcription (40). Since our model focuses specifically on polysome formation, we include only these (750-2500) transcriptionally-active RNAPs, which can transition between free and DNA-bound states.

Polysome formation follows a two-step process shown in Fig. 2a, b. In the first step, free ribosome particles sequentially assemble on DNA-bound RNAPs to form polysomes attached to the DNA, which we refer to as the DNA-bound polysomes. In the second step, the DNA-bound polysomes, upon reaching a critical size (*N*_ribo_ = 4) (41), detach from the DNA. Simulataneously with this polysome detachment, the RNAP is also released into the cytosol where it diffuses until it binds again to the DNA. We choose a high binding rate of RNAPs to DNA monomers relative to the other reactions (50 s^−1^) to ensure that most of the RNAPs in the simulation are actively engaged in the transcription. We assume each free ribosome assembles on polysomes at a constant rate *k*_on_ whether or not the polysome is bound to DNA. We specify this rate in the next section. Our LAMMPS model only attempts polysome detachment from DNA and RNAP when the number of ribosomes in the DNA-bound polysome reaches three (*N*_ribo_ = 3). After reaching this threshold number, we take the detachment rate, *k*_detach_, to be equal to *k*_on_. Once the polysome detachment attempt is accepted, *N*_ribo_ increases from three to four and the DNA-bound polysome detaches (see Fig. 2b). In this polysome formation process, the free ribosomes only attach to but do not dissociate from the DNA-bound polysomes.

**Figure 2:**
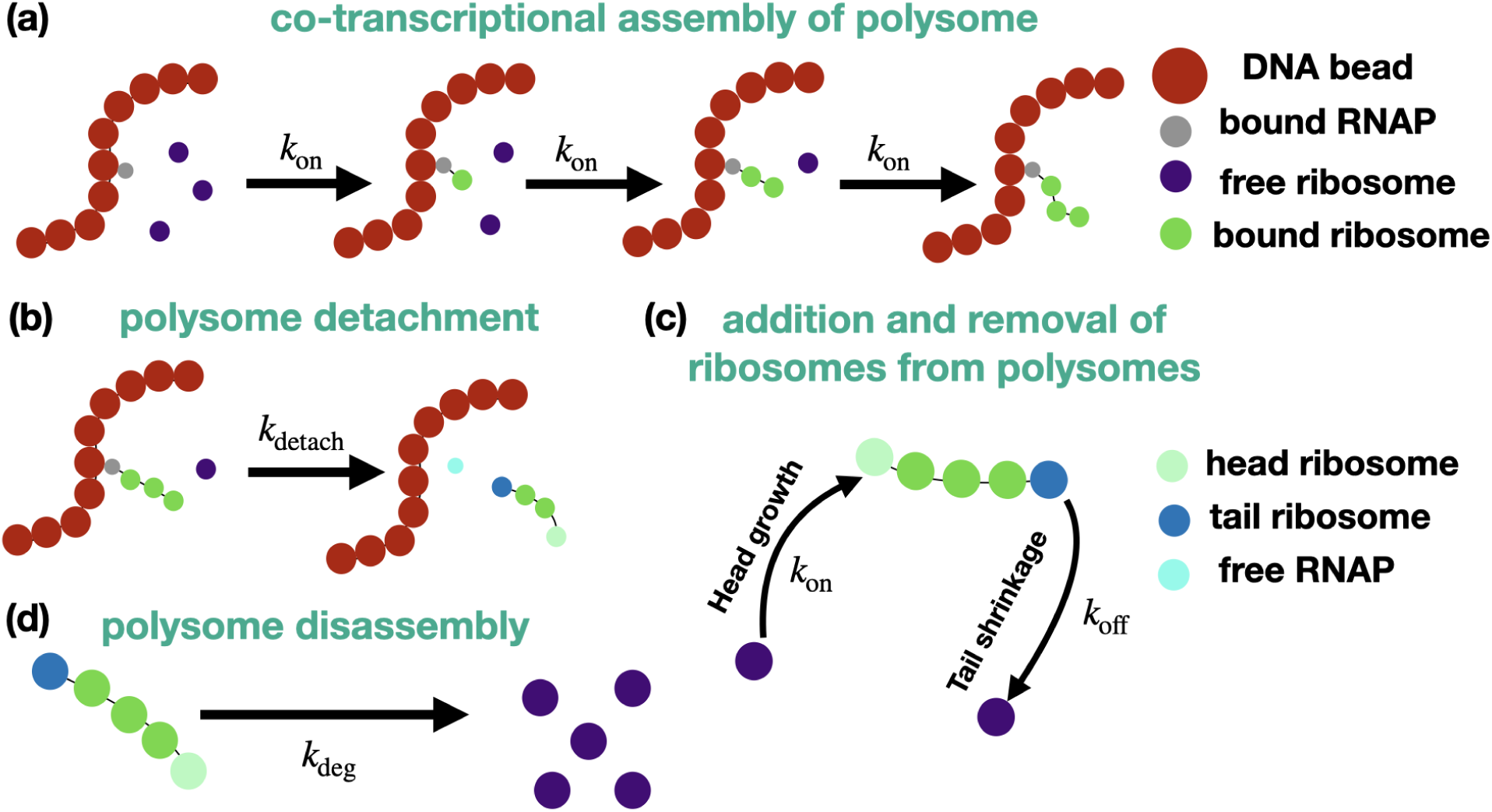
Schematics of reactions included in the model. (a) Polysome assembly initiates when a free ribosome (purple) binds to a DNA-bound RNAP (gray). This is followed by the sequential assembly of additional ribosomes (green) with rate constant *k*_on_. (b) The polysome detaches upon reaching a fixed size *N*_ribo_ = 4. Simultaneously, the RNAP also detaches and becomes a free RNAP (cyan). (c) Polysome treadmilling dynamics: head ribosomes (light green) are added with rate *k*_on_ while tail ribosomes (blue) dissociate with rate *k*_off_. (d) Polysome disassembly (the complete dissolution of the polysome complex into individual free ribosomes) occurs with rate *k*_deg_.

#### Addition and removal of ribosomes from freely diffusing polysomes

Once polysomes detach from RNAP and DNA, they grow from one end (the head) through the addition of ribosomes while shrinking from the other end (the tail) as the ribosomes complete protein synthesis and dissociate (Fig. 2c). We refer to this type of behavior as polysome treadmilling. We model the ribosome addition to the “head" of a polysome via a rate constant *k*_on_. We model the dissociation of a ribosome from the “tail" of a polysome via a rate constant *k*_off_. We estimate the value of *k*_off_ using the measured rate of translation in *E. coli* of ≈ 12 amino acids per second (39). Taking the average protein to be 350 amino acids long, the average time for a ribosome to complete protein translation is 30 seconds. Since there are, on average, 4 ribosomes per polysome (41), we expect to lose a single ribosome from a polysome every 7.5 seconds. Thus, we take the unbinding rate of ribosomes from a polysome, *k*_off_, as the inverse of this time: *k*_off_ ≈ 1*/*(7.5 s) ≈ 0.14 s^−1^.

To estimate the bimolecular ribosome binding rate *k*_on_, we consider the evolution of free ribosome concentration *c*_fr_ under well-mixed conditions where we assume the concentration *c*_fr_ is spatially uniform. In this case, its time evolution reads

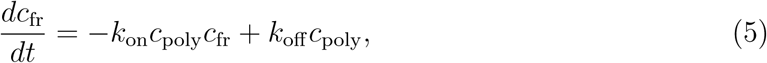

where *c*_poly_ is the (uniform) concentration of free polysomes. Assuming a steady state, we have *k*_on_ = *k*_off_ */c*_fr,steady_, where *c*_fr,steady_ is the steady-state free ribosome concentration. Previous studies report that the fraction of free ribosomal subunits in log-phase *E. coli* cells is about 20% (36, 37). *c*_fr,steady_ in our model then can be estimated to be *c*_fr,steady_ ∼ 0.2*n*_ribo_*/V*_cell_ = 1500 *µ*m^−3^, where *n*_ribo_ = 6000 is the total number of ribosomes and *V*_cell_ = 0.8 *µ*m^3^ is the cell volume. This yields *k*_on_ ≈ 9.3 × 10^−5^*µ*m^3^*/*s.

In Eq. 4 we use an effective free ribosome binding rate 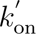 for ribosome attachment to polysome. This attachment rate is relative to a given polysome and thus is a first-order reaction. To define this first-order reaction rate, we approximate the ribosome concentration relative to the head end of the polysome as one ribosome particle in the sphere defined by the maximum reaction distance *R*^max^ = 1.1*σ*_ribo_, that is, *c*_fr_ = 3*/*(4*π*(1.1*σ*_ribo_)^3^) ≈ 2.24 × 10^5^ *µ*m^−3^. Using this approximation, we calculate 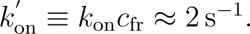

#### Polysome disassembly

Polysomes disassemble when the mRNA template is degraded, as shown in Fig. 2d. When a polysome disassembles, all FENE bonds in the polysome are removed and the ribosomes that were previously bound to the polysome are released back into the cell as free ribosome particles. The mean lifetime of mRNA in *E. coli* is reported as approximately 2.5 minutes (42). We take the corresponding degradation rate, *k*_deg_, as the inverse of this time: *k*_deg_ ≈ 6.7 × 10^−3^ s^−1^.

### Brownian thermostat

We conduct molecular dynamics (MD) simulations at fixed cell volume and temperature. The total number of ribosomal particles and DNA monomers is also fixed as the reactions considered here conserve the total number of ribosomes. The time integration is performed using the Brownian dynamics integrator in LAMMPS (25). For further details on Brownian dynamics, see APPENDIX. The length unit in our simulation is the free ribosome diameter (*σ*_ribo_ = 20 nm) and the time unit is set by the free ribosome diffusion coefficient (*D*_ribo_) via

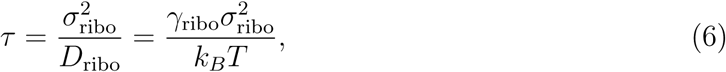

where *γ*_ribo_ is the damping constant for the free ribosomes.

We estimate *τ* from experimental data on the mean square displacement (MSD) of 25 nm nanocage-GFP particles in *E. coli* (43). We used the experimental data from nanocage particles and not from ribosome tracking experiments because the interpretation of these data is simpler. The nanocage particles, unlike ribosomal subunits, do not undergo binding and unbinding reactions. At the same time, the nanocage particles have a similar size to free 70S ribosomes.

The experimentally measured MSD vs. time curve for the nanocage particles (Fig. 3a) is non-linear. The non-linearity likely arises because of cell confinement. To verify this assumption, we build a simple LAMMPS model featuring ten ribosome particles confined to a spherocylindrical *E. coli* cell. This preliminary model does not include any reactions. To account for the macromolecular crowding in the cell, we also include in the model 600 polysomes, each consisting of *N*_ribo_ = 10 ribosomes (15, 44), and a circular DNA polymer consisting of 200 monomer units. These are the same parameters as in our non-equilibrium model described above. The MSD vs. time curve from the model shows a similar nonlinearity as the experimental data (Fig. 3a). To determine the modeling time scale *τ*, we adjust its value so that experimental and modeling MSD curved reached a reasonable agreement (Fig. 3a). We find this to occur for *τ* ≈ 5 ms. This time conversion from dimensionless modeling times to actual experimental times is used for all the subsequent LAMMPS models including those use reactions.

**Figure 3:**
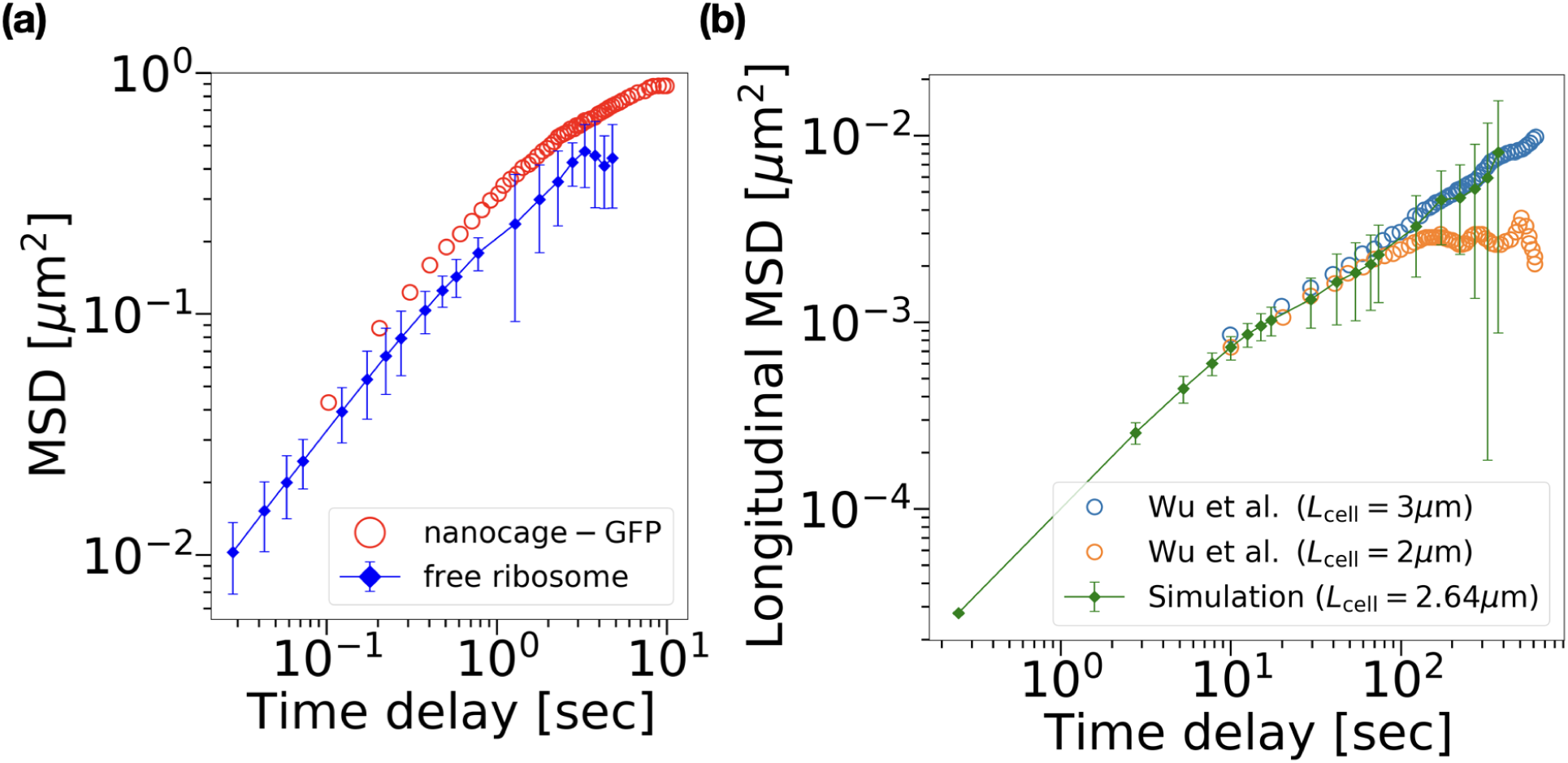
Calibration with experimental data. (a) Measured MSD of 25 nm nanocage-GFP particles from (43) (open red circle) compared with the MSD of (simulated) 20 nm diameter free ribosome particles (solid blue curve). The error bars represent the standard error of the mean over 10 different free ribosome particles in the same simulation run. (b) Longitudinal MSD of the nucleoid center of mass as a function of time: experimental data from 2 *µ*m (orange circles) and 3 *µ*m (blue circles) long cells (32) are used to compare with simulation results from ≈ 2.64 *µ*m cells (green diamonds, means ± the standard errors of mean from three independent simulations are presented).

To model the diffusion of DNA, we set the diffusion coefficient of individual DNA beads, *D*_bead_, to *D*_ribo_*/*4. We choose this value based on the Stokes-Einstein relation since the diameter of DNA monomers in our model is 4-fold larger than that of ribosomes (see Table 1). To determine the validity of this approach, we calculate the MSD vs. time curve for the center of mass (CM) of the whole chromosome and compare it to the experimental curve from Wu *et al.* (32). The model and data show a good agreement with each other (Fig. 3b) validating the approach.

We set the integration time step in LAMMPS to *dτ* = 10^−4^ *τ*. This time step is sufficiently small and avoids particle interpenetration (and resultant very high forces), which can emerge from non-equilibrium reactions. We collect the complete particle configurations every 5 ×10^5^ timesteps (50 *τ*), unless otherwise specified. The parameters used for the Brownian dynamics (single chromosome cases) are listed in Table 3.

**Table 3:** Input parameters for Brownian dynamics simulation.

|  | Symbol | Value | Unit |
| --- | --- | --- | --- |
| Diffusion coefficient of ribosomes | $D_{\text{ribo}}$ | $8 \times 10^{-2}$ (43) | $\mu\text{m}^2/\text{s}$ |
| Diffusion coefficient of DNA beads | $D_{\text{bead}}$ | $2 \times 10^{-2}$ ($D_{\text{ribo}}/4$) | $\mu\text{m}^2/\text{s}$ |
| Number of DNA beads on a chromosome | $N_{\text{bead}}$ | 200 | |
| Total number of ribosomes | $n_{\text{ribo}}$ | 6000 (39) | |
| Total number of active RNA polymerase | $n_{\text{RNAP}}$ | 750 (40) | |
| Characteristic time | $\tau$ | $5 \times 10^{-3}$ | s |
| Simulation timestep size | $d\tau$ | $5 \times 10^{-7}$ | s |

### Determining nucleoid volume

To define the nucleoid region and estimate the nucleoid volume, we apply the 3-dimensional k-d tree algorithm from the Python library *SciPy* (45) to build a point cloud representing the 3-dimensional nucleoid region. We estimate the volume of the nucleoid by first creating a 3D grid and identifying all grid points that lie within a specified distance threshold (8*σ*_ribo_) of any DNA monomer. The grid spacing is chosen to be *σ*_ribo_ (20 nm), which is the unit of length in our model. The nucleoid volume is calculated by multiplying the total number of grid points inside this boundary by the cube of the grid spacing.

### Determining nucleoid overlap

To measure the degree of the overlap between the two chromosomes (labeled as DNA_1,2_), we calculate the quantity

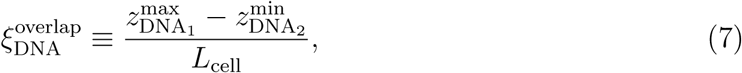

where 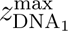 is the location, along the cell’s long axis, of the rightmost DNA monomer on the left chromosome (DNA_1_) and 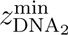 is the leftmost DNA monomer on the right chromosome (DNA_2_). Therefore, if 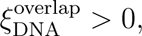 the two chromosomes overlap, and if 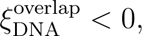 the two chromosomes are completely disentangled and separated, with a gap of 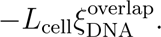

## Results

### The spatial organization of chromosomal DNA and polysomes

We first explore the effects of central dogma processes on the compaction and organization of a single nucleoid. We initialize the simulation from a state where all RNAPs are bound to chromosomes, and all ribosome particles are free; that is, there are no polysomes initially. As the simulation progresses, the majority (∼ 80%) of the free ribosome particles will assemble into polysomes (as the green and blue particles in Fig. 4a). As the system reaches a steady state, the free ribosome fraction shows only small fluctuations (∼ 4% on average, see SI Fig. S1). The distribution for the number of ribosomes in the polysome peaks at *N*_ribo_ = 4 while the average number is approximately 7 (SI Fig. S2a). The latter lies between the values that have been previously reported (*N*_ribo_ = 4 in (41), *N*_ribo_ = 10 in (44)). We calculate the MSD of the free polysomes and ribosomes in the steady state. We find free ribosomes diffuse in the same way as in the preliminary equilibrium simulation with no reactions (cf. Fig. 3a, SI Fig. S2b) that suggests the central dogma-related reactions considered here have minimal effect on the diffusivity of free ribosome particles. It is also worth noting that the polysomes diffuse on average by a factor of ten slower than free ribosomes due to their larger size in the simulation that includes reactions (SI Fig. S2b).

**Figure 4:**
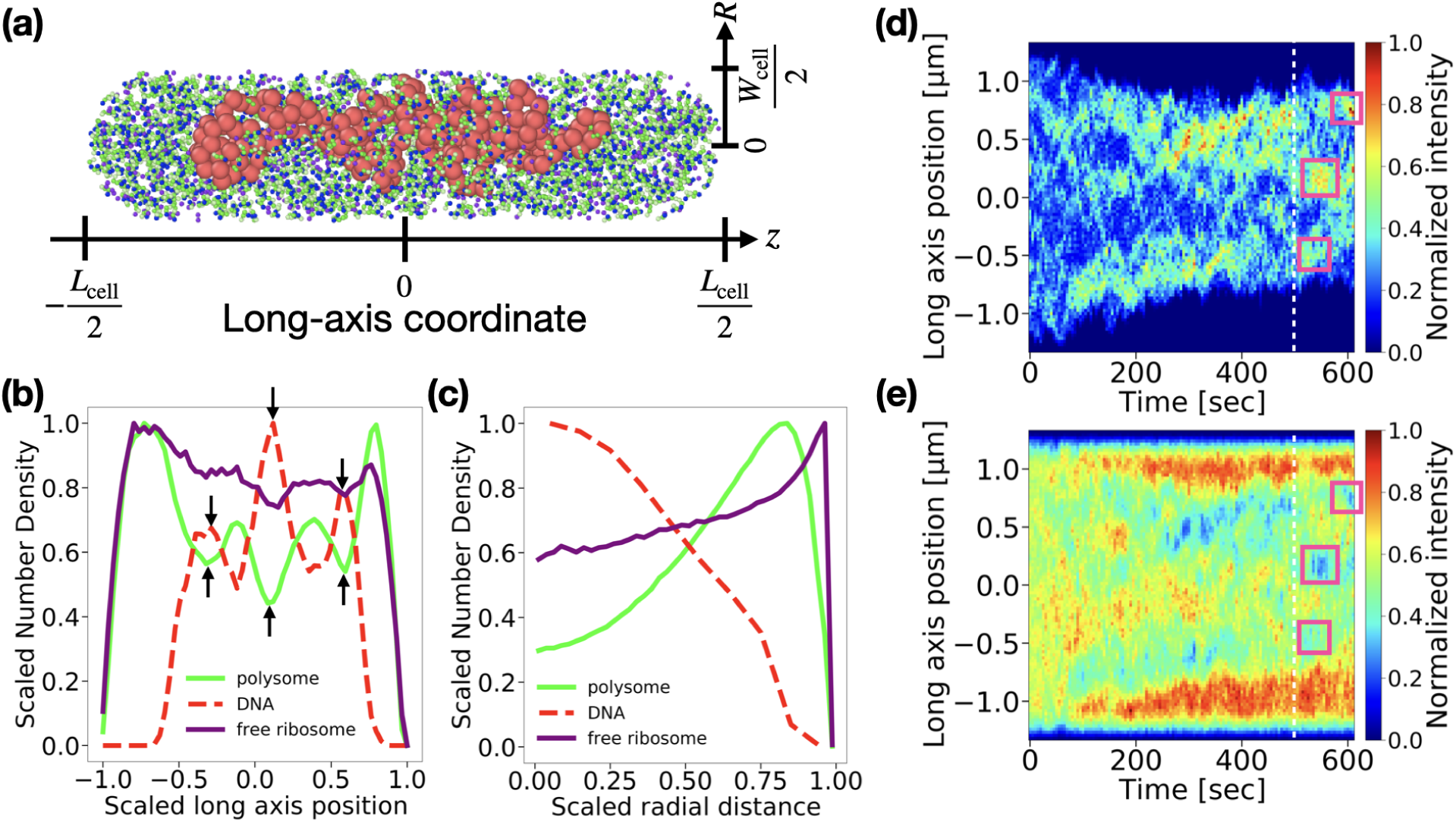
Spatial distributions of chromosome and ribosomes under out-of-equilibrium conditions. (a) A snapshot of the simulation showing DNA (red), polysomes (green and blue), and free ribosome particles (purple). For clarity, RNAPs are not shown. The coordinate system is indicated. (b) Time-averaged density distributions along the cell’s long axis for DNA (red dashed), polysomes (green), and free ribosome particles (purple). The scaled long-axis position is 2*z/L*_cell_. The arrows point to locally-anti-correlated accumulations of DNA and polysomes from a single simulation run. (c) Time-averaged radial density distributions of the mid-cell region for DNA, polysomes, and free ribosome particles from the same simulation. The mid-cell region is chosen to be from −*L*_cell_*/*4 to *L*_cell_*/*4. The scaled radial position is 2*R/W*_cell_. In both (b) and (c), the curves are averaged over the last ∼ 2 × 10^8^ steps [corresponding to ∼ 100 seconds, starting from the white-dashed line in (d)] and normalized to their respective maxima. (d,e) Kymographs showing the spatiotemporal dynamics of DNA (d) and ribosome densities (e) along the cell’s long axis from the same simulation. The white-dashed line indicates the start of the last ∼ 2 × 10^8^ steps. The magenta boxes highlight examples of local anti-correlation between high DNA density regions in (d) and the corresponding low ribosome density regions in (e).

To study the spatial distributions of species in the steady state, we collect the positions of DNA, polysomes, and ribosomes in the cell, averaging over the last ∼ 2 ×10^8^ timesteps (∼ 15% of the full simulation run). The density distributions of these species along the long-axis direction show that polysomes accumulate at cell poles and exclude DNA (Fig. 4b), consistent with previous experimental results (46). The distributions of these species along the radial direction also show apparent exclusion between DNA and polysomes, while free ribosome particles are more evenly distributed throughout the cell Fig. 4c. Similar distributions are also acquired from two additional simulations with different random seeds (SI Fig. S4). Our non-equilibrium model, like the equilibrium ones studied in earlier works (15, 44), thus shows a clear segregation of DNA and polysomes on the scale of the entire *E. coli* cell. The segregation occurs not just along the long axes of the cell but also along the radial direction. Note that previous models on crowding effects on DNA find that the DNA polymer adsorbs to the wall in the presence of crowders at equilibrium (47). To circumvent this effect some works have added a layer near the cell boundary where the crowders have been artificially excluded (4, 16). However, under the conditions of our model, which includes a significant macromolecular crowding, we find that the chromosome remains approximately in the center of the cell without any artificial layer, as is the case in live cells.

We are also interested in the organization of the DNA and polysomes on smaller spatial scales within the nucleoid. The DNA kymographs show local density fluctuations that persist for tens of seconds (Fig. 4d). Increased DNA density regions match the decreased ribosome density regions (Fig. 4d, e, marked by magenta boxes), and vice versa. The patterns suggest that local clustering of DNA excludes ribosomes, and reciprocally, ribosome-rich regions are depleted of DNA. These anti-correlated patterns are also apparent in the time-averaged DNA and density distributions (Fig. 4b, arrows). These findings are consistent with experiments indicating that DNA in the nucleoid is heterogeneously distributed and shows local clustering that anti-correlates with ribosomes (43).

### Out-of-equilibrium processes increase the size of the nucleoid

To investigate the role of out-of-equilibrium processes in nucleoid compaction, we compare simulations that include reactions to simulations in which all the reactions (as shown in Fig. 2) are stopped. We stop the reactions at the final state of the simulation shown in the previous section (the last time point in Fig. 4) and then run the simulation without reactions for about 600 more seconds (SI Fig. S4). In this latter simulation, polysomes do not dissociate, there is no ribosomal treadmilling on the phantom mRNA template, and the polysomes that are attached via RNAP to the chromosome remain attached. However, the free ribosome particles and free polysomes can move freely and redistribute in the cell.

Our findings show that the nucleoid volume *V*_nuc_ (see METHODS for details of estimating *V*_nuc_), upon deactivating reactions at *t* = 0, decreases from ∼ 0.5 to 0.47 *µ*m^3^ (∼ 6%) in 250 seconds (Fig. 5a). A similar decrease can also be observed for the radius of gyration of the chromosome along the long-axis of the cell, *R_g_*_∥_, which decreases by ≈ 20% during the same time period (Fig. 5b). Interestingly, the radius of gyration along the short (radial) axis of the cell, *R_g_*_⊥_ does not show significant change upon deactivation of reactions (Fig. 5c). An anisotropic change in the nucleoid dimension was also observed in an earlier equilibrium model when the number of macromolecular crowders in the cell increased (4). This finding from the equilibrium model was consistent with the experimental data from the same work.

**Figure 5:**
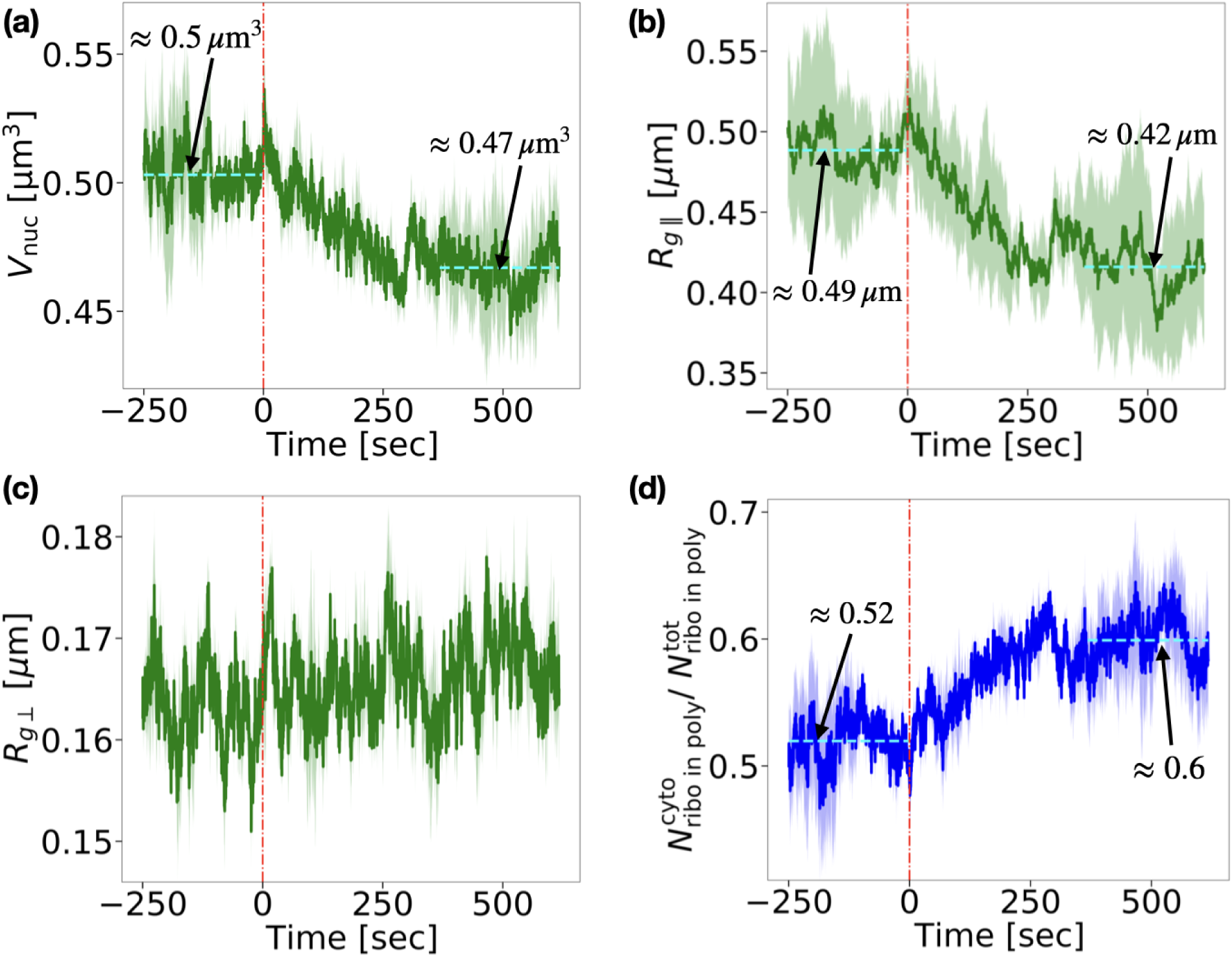
Compaction of the nucleoid after stopping the reactions. (a) Nucleoid volume *V*_nuc_ as a function of time. (b) The chromosome’s radius of gyration along the cell’s long axis *R_g_*_∥_ as a function of time. (c) The chromosome’s radius of gyration along the radial direction *R_g_*_⊥_ as a function of time. (d) 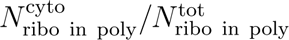 as a function of time. 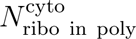 is the total number of ribosomes in polysomes in the cytosolic phase and 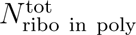 is the total number of ribosomes in polysomes in the whole cell. The horizontal cyan dashed lines in all panels indicate the average steady-state values before and after the deactivation of reactions with corresponding values shown above the lines. In all panels, the red dash-dotted line at *t* = 0 marks the deactivation of all non-equilibrium reactions. Shaded areas represent the standard error of the mean over three simulation runs.

The unchanged nucleoid dimension in the radial direction indicates that polysomes and ribosomes largely move from the nucleoid region into cell poles instead of the radial shell around the nucleoid. These excess polysomes in the cell pole regions then squeeze the nucleoid along the long axis. Consistent with this interpretation, the shrinkage of the nucleoid along the long axes of the cell is accompanied by an increase in the fraction of ribosomes in polysomes in the cytosolic region (i.e., outside the nucleoid region), 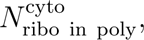 from 52% to 60% of the total number of ribosomes in polysomes in the cell, 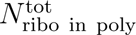 (Fig. 5d).

The data on ribosome partitioning (Fig. 5d) also shows that there is a significant number of polysomes in the nucleoid region both when the reactions are enabled (∼ 48%) and when they are stopped (∼ 40%). This high fraction differs significantly from the predictions of equilibrium models (15, 44) and from non-equilibrium continuum models (33, 35) where polysomes are almost completely excluded from the nucleoid due to their strong excluded volume interactions with DNA. A large fraction of polysomes are present in the nucleoid region in our non-equilibrium simulations because polysomes form there during co-transcriptional translation. The assembly of polysomes in the nucleoid region via co-transcriptional translation is possible because free ribosome particles are able to diffuse to the nucleoid due to their relatively low excluded volume interactions with DNA as compared to polysomes. Note that the fraction of ribosomes in polysomes that are present in the nucleoid region after reactions stop does not decay to zero in our model because DNA-bound ribosomes remain attached to DNA.

To further understand how the non-equilibrium processes affect the nucleoid compaction, we vary the number of active RNAPs while keeping all other parameters fixed in the model. We find that *R_g_*_∥_ as a function of the active RNAP number increases linearly (Fig. 6a). The linear increase results because the active RNAP numbers lead to a linear increase in the number of DNA-bound polysomes and a decrease in free ribosome numbers (Fig. 6b). The DNA-bound polysome fraction causes the size of the nucleoid to increase via the excluded volume interactions between DNA and DNA-bound polysomes.

**Figure 6:**
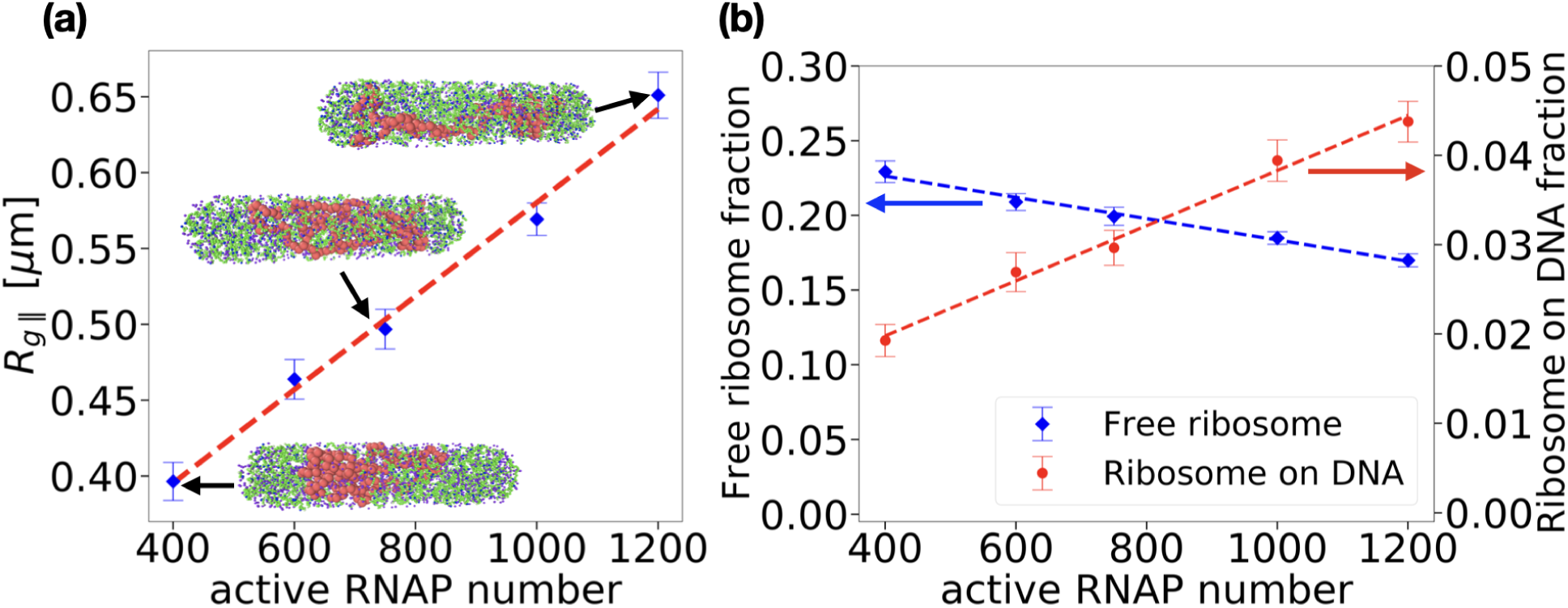
Impact of transcriptional activity on nucleoid organization and ribosome distribution. (a) The radius of gyration of the chromosome along the long axis of the cell, *R_g_*_∥_, as a function of active RNAP number. The dashed line is the linear fit to the data (*R_g_*_∥_ = 3 · 10^−4^ · *n*_RNAP_ + 0.27 and *R*^2^ = 0.992). Representative snapshots of simulations with different RNAP numbers are shown for *n*_RNAP_ = 400, 750 and 1200. (b) The fractions of free ribosome particles (blue, left axis) and ribosomes bound to DNA (red, right axis) as a function of the active RNAP number. Dashed lines are linear fits to the simulation data (*y* = −7.1 · 10^−5^ · *x* + *b, R*^2^ = 0.988 for the free ribosome fraction, and *y* = 3.1 · 10^−5^ · *x* + 0.0075*, R*^2^ = 0.990 for the ribosome fraction that are linked to DNA). The error bars in (a) and (b) represent the standard error of the mean over two simulation runs. *R_g_*_∥_ and ribosome fractions of each simulation are calculated using the last approximately 15% of the simulation steps.

### The effect of inhibiting transcription on nucleoid compaction

Next, we quantitatively compare our model to experiments where the initiation of transcription is blocked by rifampicin (Rif) antibiotic. Experiments show that the nucleoid first briefly contracts and then more slowly expands after Rif treatment in *E. coli* cells (4, 36, 43, 48). To model the effects of rifampicin, we start again from the initial condition that is defined by the endpoint of simulations in Fig. 4, representing a typical state of normally growing an *E. coli* cell in slow growth conditions before its DNA replication starts. We assume the drug is administered at time *t* = 0, at which point we stop free ribosomes from binding to bound RNA polymerase to account for the rifampicin-induced inhibition of transcription initiation. We also disable reactions involved in ribosomal treadmilling to prevent the existing polysomes from growing unrealistically long. The formation of very long polysomes would otherwise occur in our model because we do not explicitly model the finite size of mRNA, which in actual cells sets a limit on how many ribosomes can be present in a polysome. At the same time, we maintain other out-of-equilibrium reactions, including a detachment of polysomes from DNA and their dissociation (Fig. 7a). As a result of these reactions, the fraction of ribosomes linked to DNA via RNAP decreases to zero in the model, while the fraction of free ribosomes approaches 100% following the introduction of Rif (Fig. 7b). As the polysomes disassociate to free ribosomes following Rif treatment, the accumulation of ribosomes (polysomes) vanishes from the pole regions and ribosomes distribute more evenly throughout the cell (Fig. 7c right, *t >* 0). These changes in ribosome distributions are reflected in DNA density distributions that also become more uniform and spread throughout the cell (Fig. 7c left, *t >* 0).

**Figure 7:**
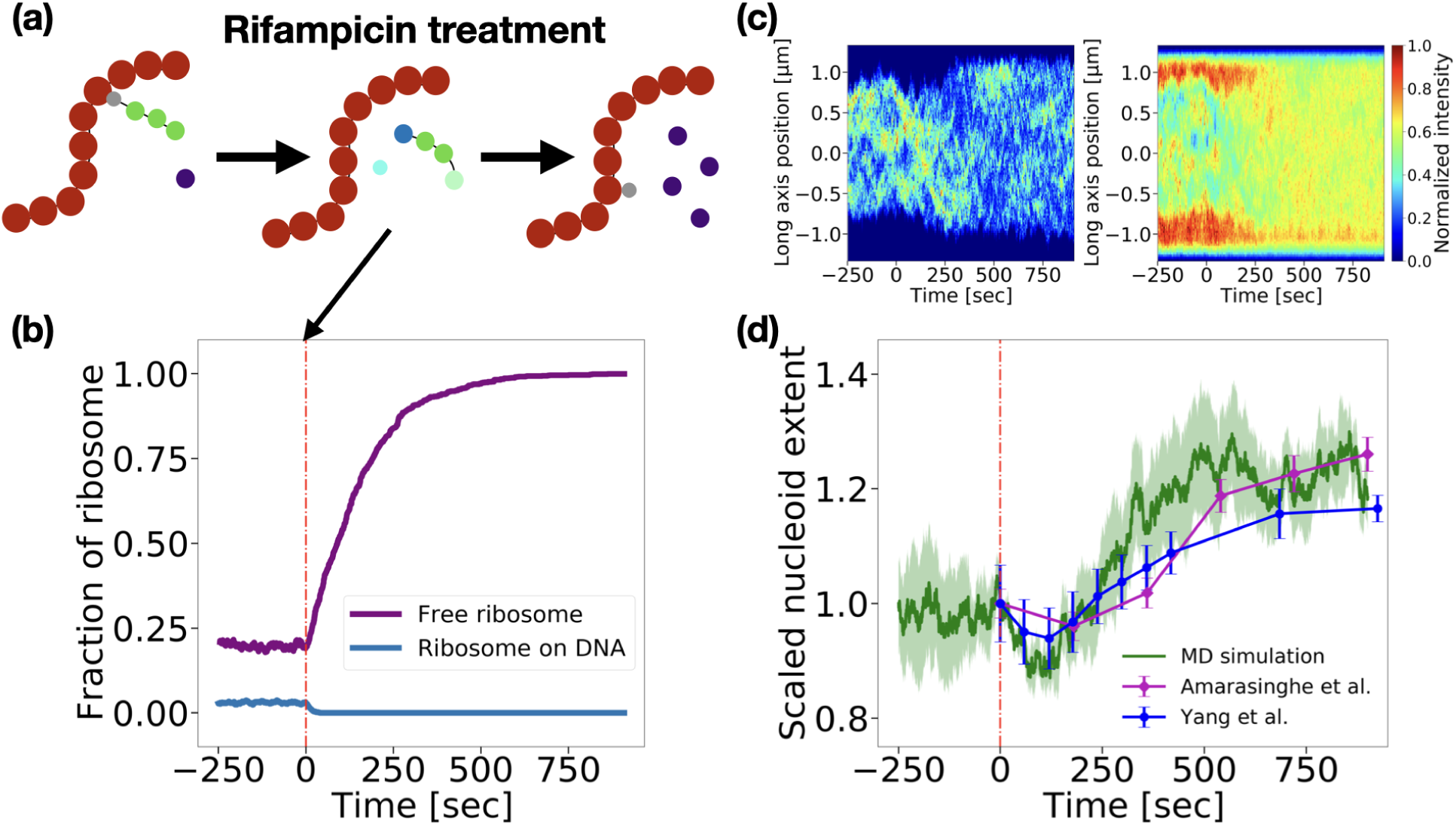
Simulation of nucleoid dynamics following transcription inhibition by rifampicin. (a) Schematic for dissociation of polysomes during Rif treatment. The ribosome-RNAP binding and polysome treadmilling are disabled, while the assembly of DNA-bound polysomes, polysome detachment and polysome disassembly are allowed to occur. (b) Time evolution of the fraction of free ribosome particles (purple) and DNA-bound ribosomes (blue) relative to total ribosomes. The red dash-dotted line at *t* = 0 marks the treatment start time. (c) Kymographs showing the distribution of DNA density (left) and ribosome density (right) along the cell’s long axis from one of the simulation runs. (d) Comparison of normalized nucleoid lengths between simulation and experimental data from two independent studies (4, 35). For simulation data (green), *R_g_*_∥_ of chromosome is normalized by *R_g_*_∥_(*t* = 0). For experimental data (purple: (35), blue: (4)), the measured nucleoid extent is normalized by the nucleoid extent at the time the rifampicin treatment reaches the cells (referred to as *t* = 0). The shaded area represents the standard error of the mean over three simulation runs. Error bars for the experimental data represent also the standard error of the mean over cell population.

To test the model, we compare it to experimental measurements of nucleoid lengths in Rif-treated cells in slow growth conditions (4, 35). The cells in these measurements are in the early stages of the cell cycle containing mostly a single fully replicated DNA molecule. The nucleoid length in both modeling and experiments shows initial contraction lasting approximately 100 seconds, followed by significant expansion after about 150 seconds from the start of the treatment (Fig. 7d). The initial contraction matches approximately the compaction observed in our earlier simulations where all reactions were turned off at time *t* = 0 (Fig. 5a, SI Fig. S5). This compaction corresponds to free polysomes (polysomes not bound to DNA) diffusing out from the nucleoid region as explained before. The compaction phase is overtaken by expansion (*t* ≳ 150 seconds) when most polysomes have dissociated to free ribosome particles (cf. Fig. 7b and Fig. 7d). The excluded volume interactions between the free ribosomes and DNA are much weaker than the excluded volume interactions between the corresponding number of polysomes and DNA, leading to the expansion of the nucleoid. The same explanation for the nucleoid expansion has also been proposed based on equilibrium models (15, 44). However, in equilibrium models, polysomes disassemble instantaneously and the nucleoid size adapts instantly to the new conditions. Conversely, in our LAMMPS model, we observe the dynamic changes in both the nucleoid extent and the polysome distribution. The time scale of these changes in our model matches the experimentally observed time scale.

The initial contraction of the nucleoid in the previous experimental works has been interpreted as a consequence of the breakage of transertional linkages between the DNA and the cell’s inner membrane (4, 48). The transertional linkages are transient structures that consist of DNA-attached RNAP complexes, nascent mRNA being transcribed, and cotranscriptionally attached ribosomes (polysomes) that have started protein translation. The linkages also include translocon complexes (e.g., SecYEG) to which the N-terminus signal peptide of the protein that has been partially translated has already been inserted (10). By connecting the DNA to the cell’s inner membrane, the transertional linkages are expected to expand the nucleoid (10, 49). Rif stops the initiation of transcription so that new linkages cannot form while the existing ones vanish as the transcript is completed. As the old linkages fall off, we thus expect the nucleoid to contract. While this is a plausible scenario, our modeling shows that initially observed nucleoid compaction after Rif treatment can originate from a different process. Instead of losing the membrane-connecting transertional linkages, our model shows that losing only DNA-attached co-transcriptionally formed polysomes is sufficient to shrink the nucleoid. The explanation offered by our model is thus simpler than the one based on transertional linkages.

### Separation of two sister chromosomes

We now move to the second question of this study - how do active processes affect the separation of two daughter chromosomes from each other once they are fully replicated? We extend the previously described model by adding a second circular DNA molecule and doubling the number of RNAPs and the total number of ribosomes. We also double the cell volume by increasing the cell length (from 2.64 to 5.04 *µ*m) while keeping the cell width fixed. We start the simulations from an initial configuration with the two chromosomes completely overlapping, and all ribosomes are present as free ribosomes distributed uniformly throughout the cell and RNAPs attached to DNA (SI Fig. S6a). As the simulations progress from this initial state, the free ribosome particles assemble to form polysomes and the composition of ribosome species reaches a steady state as in the one chromosome simulations (recall Fig. S1).

At the beginning of the simulations, the overlapping chromosomes segregate via the entropic mechanism (Fig. 8a left and SI Fig. S6b) as expected from previous works (27, 28, 31). This initial segregation process is not of interest here because our model does not take into account the concurrency between DNA replication and segregation and the correct topology of DNA during the replication stage. Instead, we use this portion of the simulation to create an initial condition for a further study of chromosome separation. At the end of the segregation process, we define a reference time *t* = 0 when only a small portion of the chromosomes are overlapping 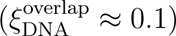 (Fig. 8a right). Starting from this configuration, we conducted independent simulations with different random seeds to understand how the nucleoids separate from this point on in the presence and absence of reactions.

**Figure 8:**
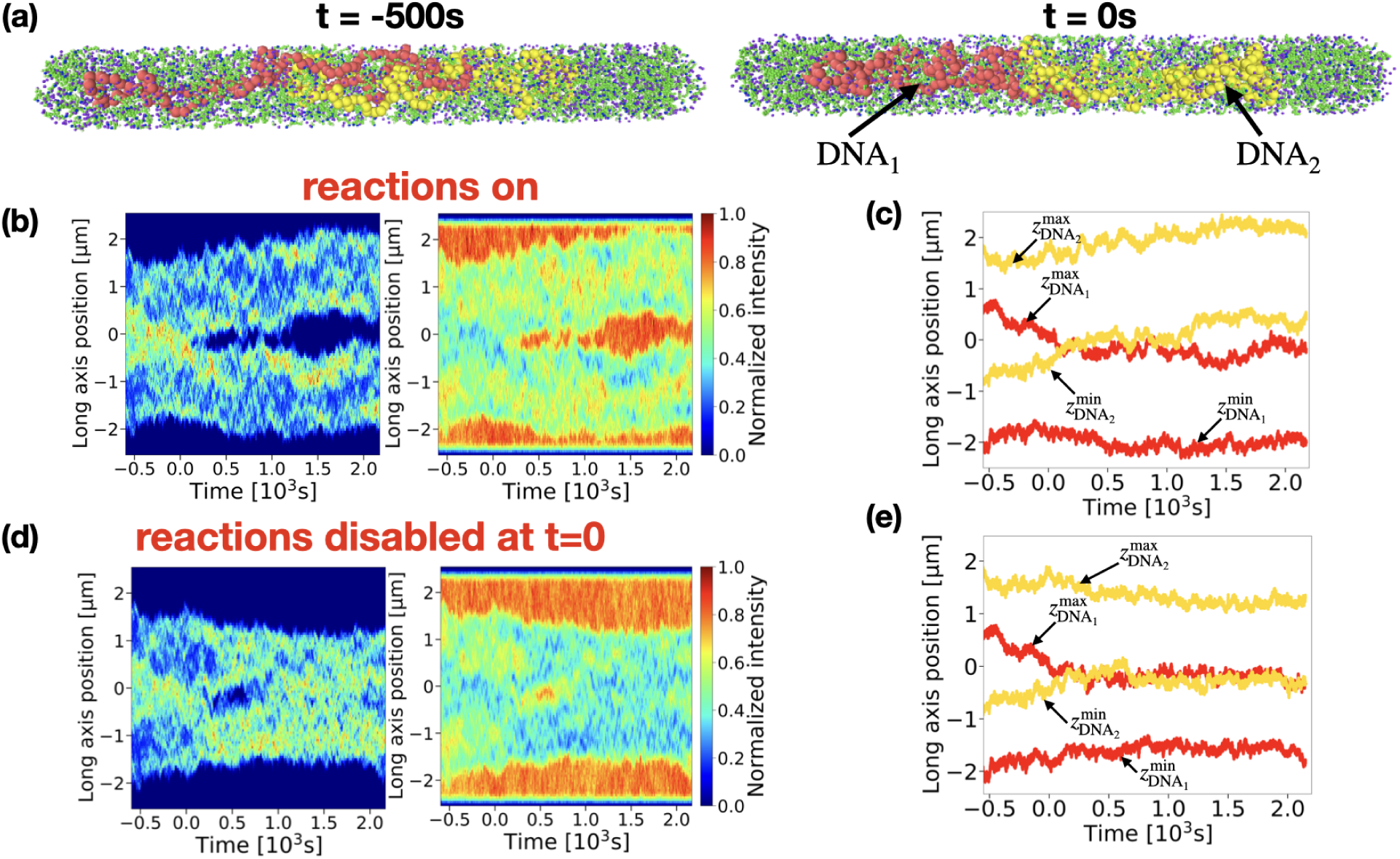
The effects of out-of-equilibrium reactions on the separation dynamics of two daughter chromosomes. (a) Snapshot of the configurations with two partially overlapping chromosomes at *t* = −500 second (left) and *t* = 0 second (right). Starting from the configuration at *t* = 0, two different sets of simulations have been conducted. One set keeps all the parameters unchanged (with out-of-equilibrium processes active), while all out-of-equilibrium processes are disabled in the other set. (b) Kymographs of the DNA density (left) and ribosome density (right) along the cell’s long axis from a single representative simulation with two chromosomes, where all out-of-equilibrium reactions are active. (c) Axial positions of the outermost DNA monomers as a function of time from the same simulation. Red curves indicate the two outermost monomers of one DNA (DNA_1_), and yellow curves of another DNA (DNA_2_). (d) Kymographs of the DNA (left) and ribosome densities (right) along the cell’s long axis from a simulation in which all out-of-equilibrium processes are disabled at *t* = 0. (e) Axial positions of the outermost DNA monomers as a function of time from this simulation.

In the presence of reactions, the two nucleoids separate from each other, forming a gap between them 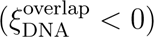 where DNA density reaches zero (Fig. 8b left and SI Fig. S7a, d). This gap is populated by polysomes (Fig. 8b right and SI Fig. S7b, e). The gap fluctuates initially between opening and closing, but eventually a persistent gap develops in all three simulations (for *t >* 1000 seconds). To quantify this behavior, we plot the positions of the outermost DNA monomers for each nucleoid on the long axes of the cell (Fig. 8c) and calculate the overlap fraction of the monomers (Fig. 9a, green curve. See METHODS for details of estimating the overlap fraction).

**Figure 9:**
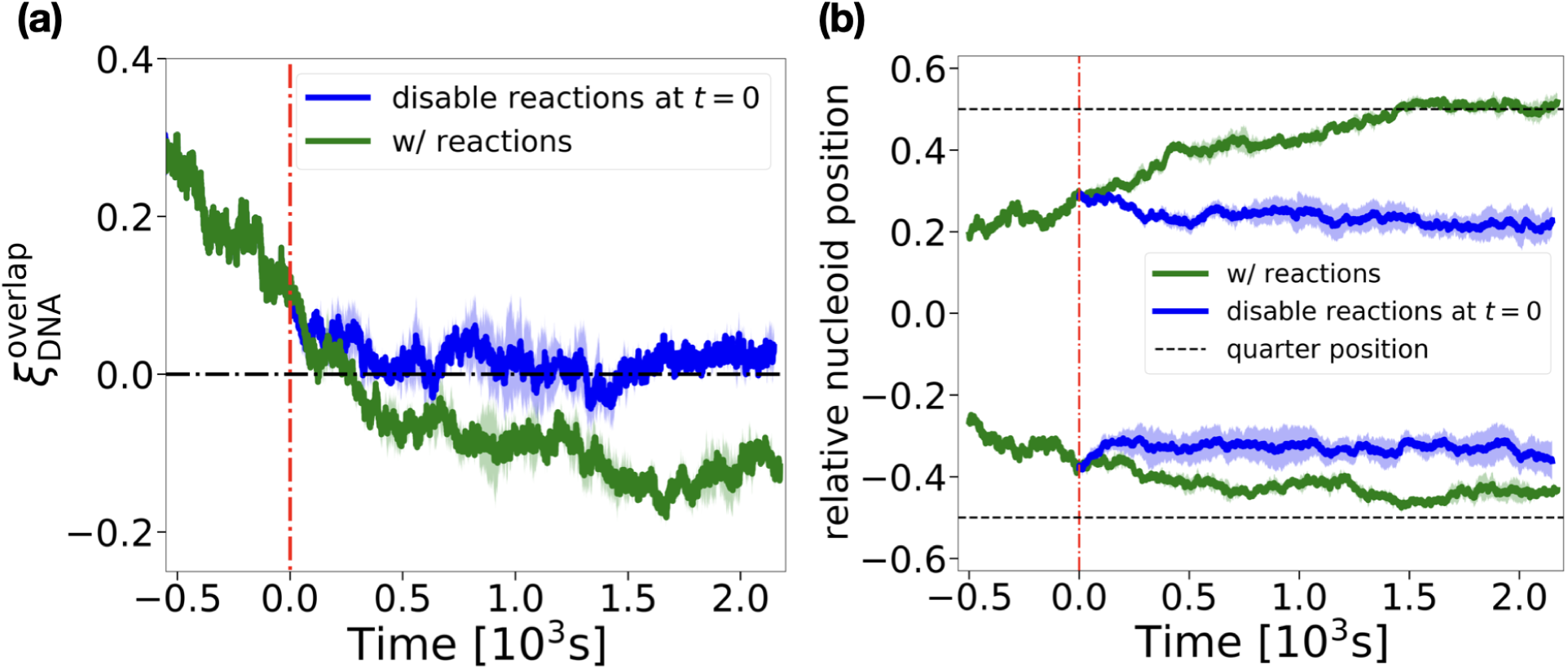
Comparison of nucleoid separation dynamics with and without reactions. (a) The time-evolution of the nucleoid overlap 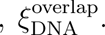 The red dashed line marks *t* = 0. The black dash-dotted line marks 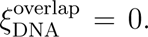 The green curve represents the simulation runs with all out-of-equilibrium reactions present, while for the blue curve, all reactions are stopped at *t* = 0. (b) The time evolution of nucleoid center of mass positions. The positions are normalized by the half of cell length *L*_cell_*/*2 = 2.52 *µ*m, such that the quarter positions correspond to 0.5 and −0.5. The shaded areas represent the standard error of the mean over three simulation runs.

We compare this behavior with simulations where all reactions are disabled from time *t* = 0. These simulations start from the same initial configuration (Fig. 8a right). The results of this simulation set show that, without the reactions, the two chromosomes do not completely separate, although transient gaps between them can occur (Fig. 8d and SI Fig. S8). In these equilibrium simulations, the only driving force that can separate the two chromosomes is the entropic repulsion mechanism (27, 50). However, the entropic force vanishes once the two chromosomes do not overlap anymore (51). At the same time, macromolecular crowders (polysomes and free ribosomes) do not distinguish one DNA molecule from another and, therefore, tend to push the chromosomes together so that diffusion of two nucleoids away from each other is hindered. However, the crowder-induced compaction of nucleoids does not yield their significant overlap in our model because it is opposed by the entropic repulsion of DNA molecules. As a result of these two opposing processes, the two nucleoids stay nearly touching with almost a zero overlap and with occasional transient gaps forming between them (Fig. 8e and Fig. 9a, blue curve).

The comparison of equilibrium and non-equilibrium simulations shows that the central dogma reactions actively drive the separation of sister chromosomes. Without these reactions, a complete separation of the two sister chromosomes would not occur. In this respect, the model has qualitatively similar behavior to the 1D continuum models (33, 34, 35), although these continuum models do not capture the fluctuations and the initial segregation process. Notably, our simulations predict that the separation between the nucleoids (*ξ*^overlap^) in the presence of reactions shows large fluctuations with the gap between the two DNA molecules opening and closing repeatedly over more than 10 min period (Fig. 8b and SI Fig. S7). This finding is in contrast to a continuum model where the separation is deterministic and abrupt, occurring in less than a minute time period (35). The fluctuating opening of the nucleoid gap in our MD model appears to be in qualitative agreement with the experimental data (35).

Experiments with filamentous cells (cell length ranges from ∼ 10 to 30 *µ*m) with two nucleoids showed that the centers of these nucleoids positioned robustly at the cell’s quarter positions, leaving a large DNA-free gap (up to tens of *µ*m) between them (32). This positioning pattern was reproduced in the model by Miangalorra *et al.* (33). The similar positioning pattern were also observed for sister chromosomes in normal *E. coli* cells (cell length ranges from 2 to 5 *µ*m) (52, 53). To see if the latter behavior could be reproduced by our model, we plot the center of mass of each nucleoid as a function of time (Fig. 9b). We find that the nucleoid centers indeed gradually approach cell quarter positions over about time *t* ∼ 1500 s. In contrast, in simulations where the reactions were stopped at time *t* = 0, the centers of nucleoid do not move further apart and remain closer to the cell center. These findings suggest that out-of-equilibrium processes are needed for achieving the experimentally observed chromosomal positioning in *E. coli* cells with two nucleoids.

## Discussion and Conclusion

In this study, we develop a Brownian dynamics model that includes out-of-equilibrium central dogma reactions. We use this model to understand the compaction of nucleoids and the separation of two chromosomes in a spherocylindrical bacterial cell such as *E. coli*. Previous molecular dynamics simulations addressing these processes have relied on equilibrium models that did not include reactions. Moreover, earlier simulations did not consider time-dependent effects. Our coarse-grained model allows us to simulate the cell over many minutes and compare model predictions to experimental data on nucleoid expansion in rifampicin-treated cells and to other time-dependent phenomena.

Our molecular dynamics simulations show that out-of-equilibrium reactions lead to larger nucleoids. This finding is consistent with existing qualitative experimental observations. It has been observed that in poorer quality medium (slow-growth conditions), the nucleoids from *E. coli* cells appear “diffuse” in microscopy images (40, 54). This broadened appearance has been interpreted to arise from high transcriptional activity throughout the genome as cells need to express transcripts for numerous enzymes involved in amino acid synthesis. Conversely, in a high-quality medium (fast-growth conditions), the transcriptional activity becomes concentrated in ribosomal operons (11) which do not lead to polysome assembly. At the same time, the cell decreases overall transcriptional activity because it does not need to express genes for amino acid synthesis. In these fast-growth conditions, *E. coli* nucleoids appear more structured and feature individualized clusters (54). While our model does not yet account for the high transcriptional activity at ribosomal operons, it predicts that regions of the chromosome with lower transcriptional activity will be more compacted, consistent with a more structured nucleoid in fast-growth conditions.

In the second part of this study, we addressed the question of how two daughter chromosomes separate from each other as they partition into two daughter cells. It has been proposed that an active mRNA-ribosome dynamics drives this process (33) in filamentous *E. coli* cells. The recent experiments combined with further modeling provide evidence that this mechanism is also important in regular *E. coli* (34, 35). However, the underlying idea is based on the results from 1D reaction-diffusion models (33, 34, 35) that do not include fluctuations and rely on virial expansions to take into account excluded volume interactions. Reducing the cell to only one dimension in these models means that the nucleoid uniformly occupies the whole cross-section of the cell. However, experiments (36) and results from our current 3D model show [Fig. 4(c)] that there is a layer of ribosomes (polysomes) surrounding the nucleoid in the radial direction of the cell. This shell-like layer was added *ad hoc* (empirically) to one of the previous models (35) but without a solid justification. It is conceivable that there is a much smaller effective force from the active mRNA-ribosome dynamics in separating the daughter nucleoids if the polysomes can diffuse away from the mid-cell region via the outer shell region (33). Although our 3D model shows the presence of the outer shell of ribosomes, it also shows that the active mRNA-ribosome dynamics is needed to separate the nucleoids. Without the reactions, the two nucleoids remain in contact in our model. The reason why the osmotic pressure on the two daughter nucleoids from the mid-cell (mid-nucleoid) polysomes is not shunted by the outer layer in our model is likely due to the slow diffusion of the polysomes. The low mobility of polysomes prevents the polysome concentrations between the mid-cell and polar regions of the cell equilibrate via the polysome diffusion through the outer shell region. Due to this persistent concentration difference, there is an outward osmotic pressure on the nucleoid from the mid-cell region to the poles which can separate the chromosomes, consistent with recent experiments (34, 35). These arguments imply that low diffusivity of polysomes is required for their dynamics to drive nucleoid separation.

The 1D reaction-diffusion model used in previous works treated the DNA as a scalar concentration field, with interaction terms determined by the excluded volume of unconnected cylinders, representing DNA segments (33, 34). The DNA concentration field can form two or more disjoint blobs even when there is only a single chromosomal amount of DNA in the model, provided the concentration of ribosomes is sufficiently high. For chromosomal DNA in cells, this outcome is prevented due to chain connectivity. In our 3D single-chromosome model, no DNA density minimum formed at mid-cell (see Fig. 4). The computational cost of the model prevented us from carrying out a series of simulations with higher ribosome concentrations, but it is clear that the three-dimensional model would predict a significant DNA density at mid-nucleoid due to the chain connectivity even at higher ribosome concentrations. The PDE-based reaction-diffusion equations are thus unlikely to correctly predict the kinetics of the formation of the DNA density minimum at mid-cell due to the lack of chain connectivity in the description. Indeed, a recent comparison of experiments and PDE-based models shows that the experimental kinetics of this process is more than an order of magnitude slower than predicted by the model (35). The cell-growth-dependent kinetics has not been addressed yet in our 3D model because it would require significantly more computational time, but it could be incorporated in future studies.

Our MD simulations provide several new predictions that could be experimentally verified. First, we find that transcriptional activity expands the nucleoid and that the expansion is proportional to the number of active RNAP in the cell. This prediction can be verified by determining nucleoid dimensions in cells where transcriptional activity is partially inhibited by a sub-lethal treatment by rifampicin or by some other means. The nucleoid dimensions can also be measured in conditions where the number of RNAP is either up- or down-regulated. The difficulty in these measurements is that transcriptional activity can also control the levels of macromolecular crowders in the cell. These levels need to be independently determined.

Secondly, our model predicts that a complete inhibition of transcription by a high concentration of rifampicin causes an initial compaction of the nucleoid followed by expansion. This behavior has already been observed in experiments (4, 48). However, the previous interpretation of the initial contraction has been that rifampicin treatment severs so-called transertional linkages that connect chromosomal DNA to the cell membrane. Our modeling suggests a simpler mechanism that can explain this experimental observation — instead of disruption of transertional linkages, the contraction may arise simply from the cessation of co-transcriptional translation and dissociation of nascent polyribosomes from DNA. Further experiments are needed to test the validity of one versus the other scenario.

Finally, the nucleoid segregation and separation dynamics warrant further experimental investigations. The existing experimental evidence shows that active mRNA-ribosome dynamics is not the sole mechanism that enables the daughter nucleoids to separate from each other as they partition into two new daughter cells (34, 35). It will be valuable to compare our existing model quantitatively with these experimental data to further validate the interpretation of these experiments, as well as test the validity of the current model.

The current model is the first attempt of a molecular dynamics that incorporates the non-equilibrium central dogma processes to understand nucleoid organization and dynamics in bacterial cells. There are many avenues to improve the current model. This includes considering more complicated DNA topologies corresponding to partially replicated chromosomes and incorporating cell growth and DNA replication in the model. It would also be advantageous to reduce the coarse-graininess of the DNA representation by using smaller DNA monomers. This would enable one to define different transcriptionally active regions within the chromosome as observed in Hi-C data (55). With smaller monomer sizes on the order of tens of base pairs, the effects arising from DNA supercoiling and looping by SMC proteins could also be investigated within the same framework. The finer-grained model could also more realistically simulate the transertion process (10), which may impact the size and structure of the nucleoid. Our model also allows us to probe fluctuations occuring both due to the thermal motion of the molecules and the stochastic nature of the implemented reactions. A detailed study of these fluctuations and a comparison to experiments is an avenue for future research.

In conclusion, our study demonstrates that bacterial chromosome organization and dynamics cannot be fully understood through equilibrium physics alone. The energy-driven central dogma processes expand the nucleoid and enable daughter chromosomes to separate from each other. We hope these new predictions from the model will be tested in future experiments.

## Acknowledgments

The authors thank Steve Abel, Chathuddasie Amarasinghe, Jaana Männik and Yuqing Qiu for useful discussions. We also thank Da Yang for help in experiments. Molecular dynamic simulations research was supported by the Center for Nanophase Materials Sciences (CNMS), which is a US Department of Energy, Office of Science User Facility at Oak Ridge National Laboratory. This research used computational resources of the Oak Ridge Leadership Computing Facility (OLCF) at the Oak Ridge National Laboratory, which is supported by the Office of Science of the U.S. Department of Energy under Contract DE-AC05-00OR22725. This work was supported by the National Institutes of Health award GM127413 (JM) and NSF MCB2313719 (JM).

## Appendix

## Brownian dynamics

The Brownian equation of motion assumes that inertial forces are negligible, as can be expected inside the viscous, crowded environment of the *E. coli* cell. The stochastic equation of motion for the position **r***_i_* of particle *i* is given by

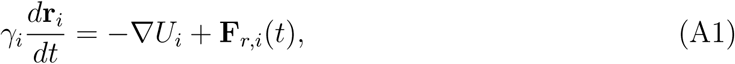

where **r***_i_* is the position of the particle, *γ_i_* is the damping constant, and **F***_r,i_*(*t*) is the fluctuating force exerted on the particle by the solvent. The random force **F***_r,i_*(*t*) satisfies the fluctuation dissipation theorem such that ⟨**F***_r,i_*(*t*)⟩ = 0 and ⟨**F***_r,i_*(*t*_1_) · **F***_r,i_*(*t*_2_)⟩ = 6*γ_i_k_B_Tδ*(*t*_1_ − *t*_2_), with *δ*(*. . .*) the Dirac delta function. The potential *U_i_* is the sum of all the interaction potentials associated with the particle *i*. For spherical particle species *i*, the damping constant *γ_i_* is given by the Stokes-Einstein equation (56):

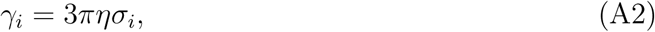

where *η* is the dynamic viscosity, assumed constant. The equation of motion in Eq. A1 is integrated using the forward Euler method:

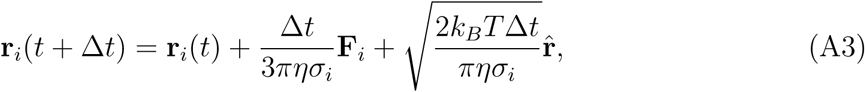

where **F***_i_* = −∇*U_i_*. In the LJ unit system, we measure the time in units of *τ*, the distance in units of *σ*_ribo_, and the energy in units of *ɛ* = *k_B_T*. We can use Eq. A3 to relate the time constant to the damping and diffusion coefficients as

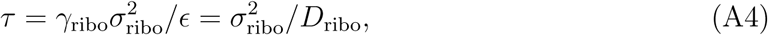

where *D*_ribo_ = *ɛ/γ*_ribo_ is the effective ribosome diffusion coefficient.

