## Supplemental Information for "Effects of central dogma processes on the compaction and segregation of bacterial nucleoids"

### SI Figures

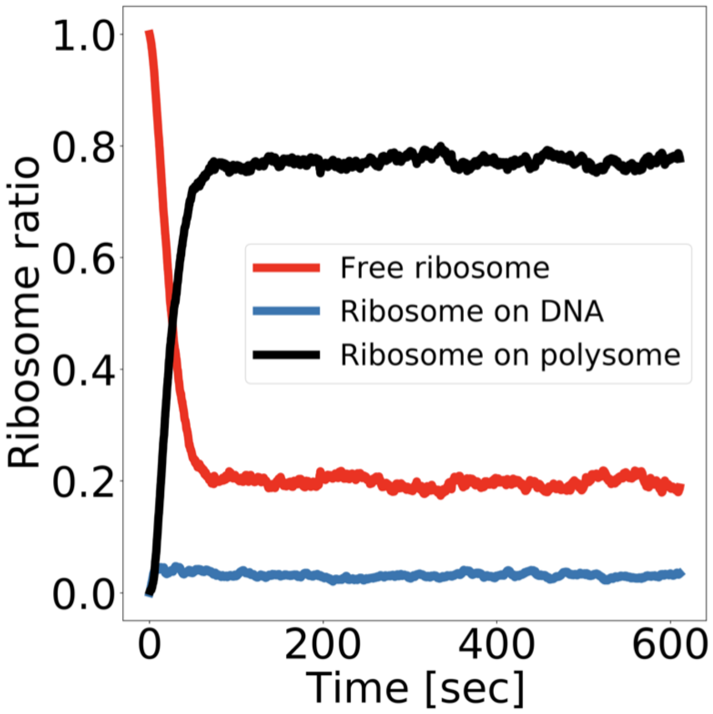


Figure S1: **The temporal evolution of ribosome species in the early stage of simulations.** The fractions of the free ribosome species (red), ribosomes bound to DNA via RNAPs (blue), and ribosomes on freely diffusive polysome (black) among all types of ribosomes in the simulation are shown as functions of time. Initially, all ribosomes are free. The free ribosome fraction then rapidly decreases as the ribosomes bind to DNA-bound RNAPs and form polysomes. The distribution of ribosome species reaches a steady state after approximately 70 seconds, where about 20% of ribosomes remain free (fluctuation $\sim$ 4% on average), while a small fraction (approximately 3%) are directly associated with DNA-bound RNAPs.

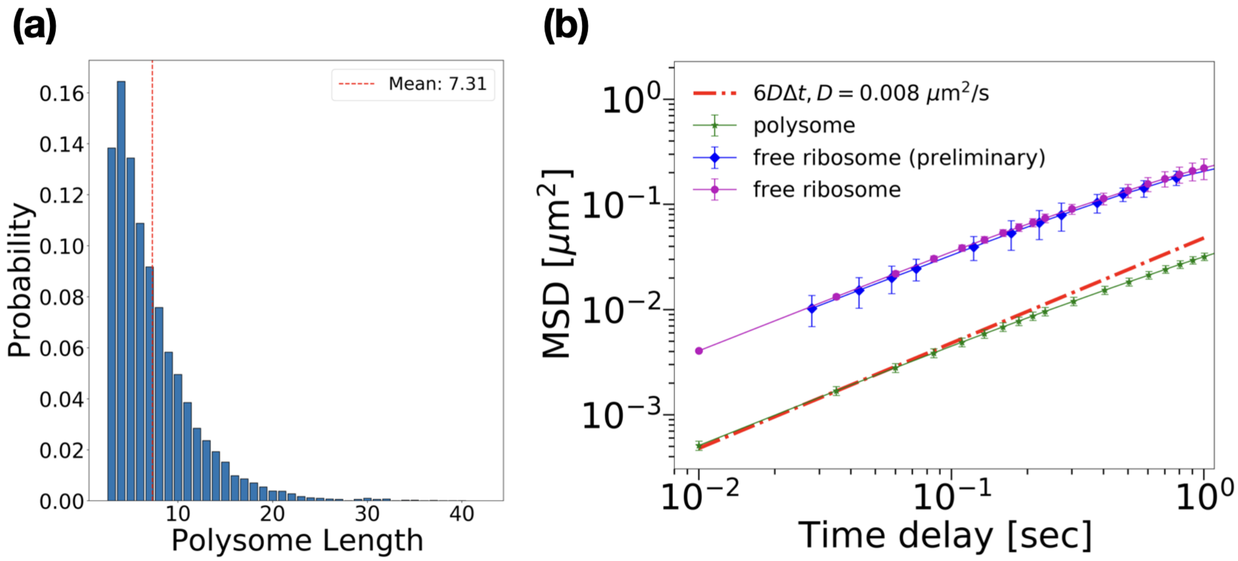


Figure S2: **The length distribution and diffusivity of free polysomes in steady state conditions.** (a) Distribution of the number of ribosomes in free polysomes ($N_{\mathrm{ribo}}$) in the out-of-equilibrium simulations. The distribution is extracted from the last $\sim 2 \times 10^8$ timesteps of the simulations ($\sim$ 100 seconds). The red-dashed line shows the average length of the polysomes in units of number of ribosomes. (b) The three-dimensional mean square displacement (MSD) of the center of mass of 10 randomly selected polysomes are presented as a function of time delay $\Delta t$ (green). The red dash-dotted line represents MSD $= 6D\Delta t$, where the diffusion coefficient is chosen as $D = 0.008\ \mu\mathrm{m}^2/\mathrm{s}$ to fit the data from simulations. The magenta curve shows the MSD of free ribosomes from the same simulation while the blue curve represents the MSD of free ribosomes from a preliminary equilibrium simulation where each polysome consists of a fixed number $N_{\mathrm{ribo}} = 10$ ribosomes (the same curve as shown in Fig. 3(a) of the main text).

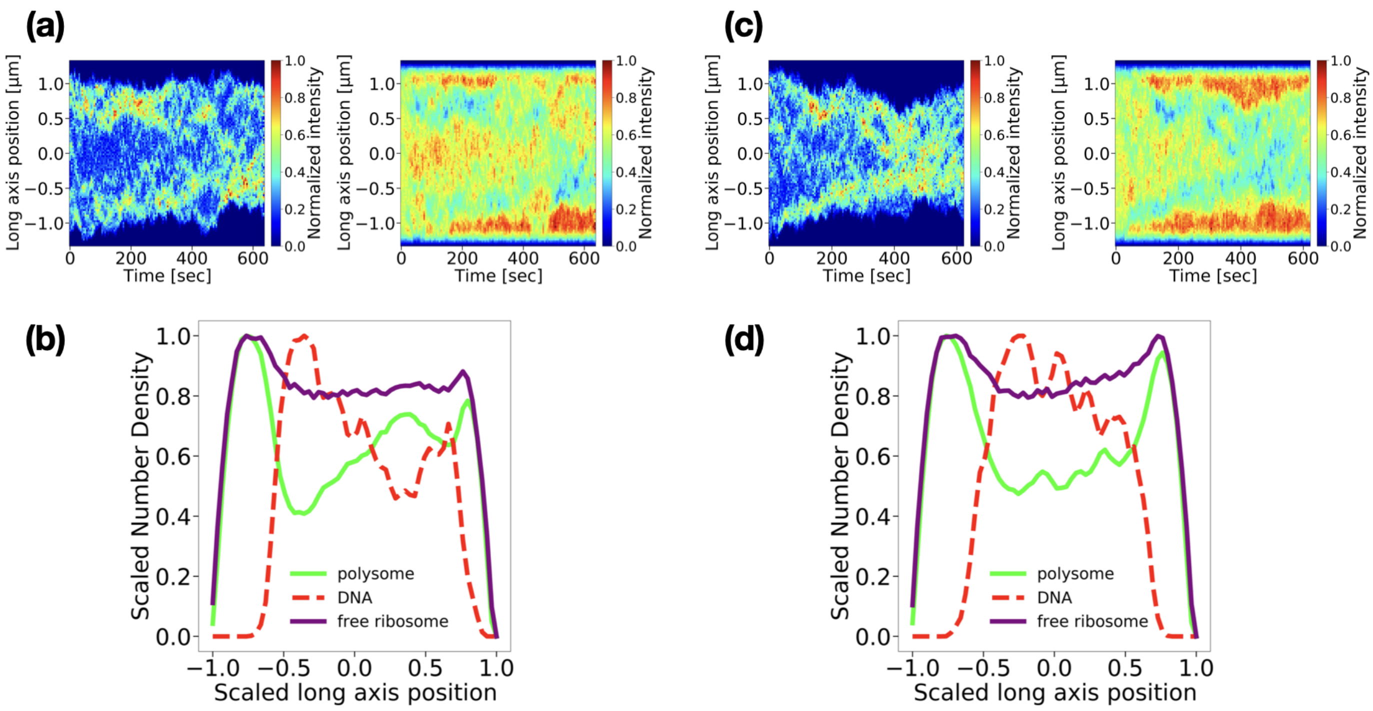


Figure S3: **Spatiotemporal distributions of the chromosome and ribosomes under out-of-equilibrium conditions from two additional simulation runs.** The two simulation runs have exactly the same parameters as the one in Fig.4 of the main text but different seed. (a, c) Kymographs showing the spatiotemporal dynamics of DNA (left) and ribosome densities (right) along the cell's long axis. (b, d) Time-averaged density distributions along the cell's long axis for DNA (red dashed), polysomes (green), and free ribosome particles (purple). Scaled long-axis position $\equiv 2z/L_{\text{cell}}$ as illustrated in Fig. 4 of the main text. The values of number density are averaged over the last $\sim 2 \times 10^8$ steps ($\sim$ 100 seconds) and normalized to their respective maxima.

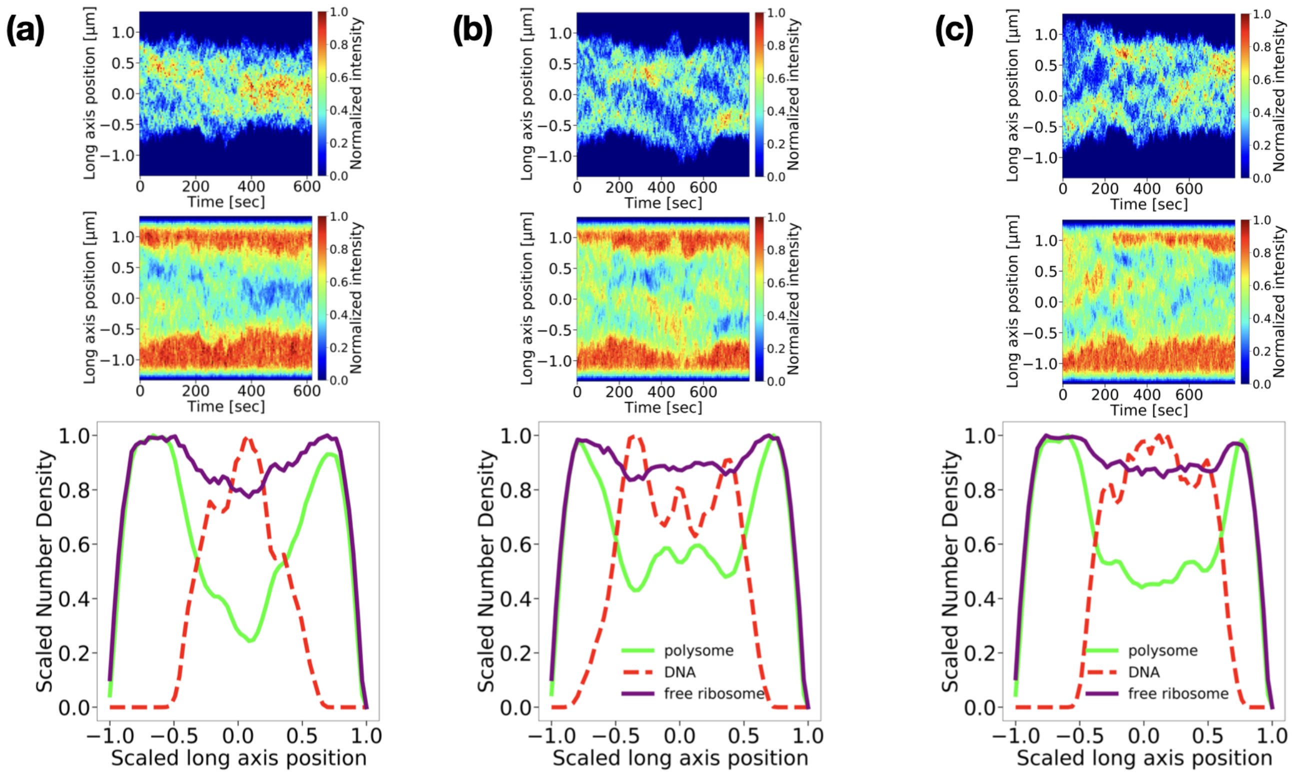


Figure S4: **Spatiotemporal distributions of the chromosome and ribosomes under equilibrium conditions (reactions disabled).** (a), (b), and (c) show the Kymographs of DNA (top) and ribosome densities (middle) and time-averaged density distributions (bottom) along the cell's long axis from three independent simulations, respectively. In the time-averaged density distributions, scaled long-axis position $\equiv 2z/L_{\text{cell}}$ as illustrated in Fig. 4 of the main text is used. The values of number density of DNA (red dashed), polysomes (green), and free ribosome particles (purple) are averaged over the last $\sim 2\times10^8$ steps ($\sim 100$ seconds) and normalized to their respective maxima.

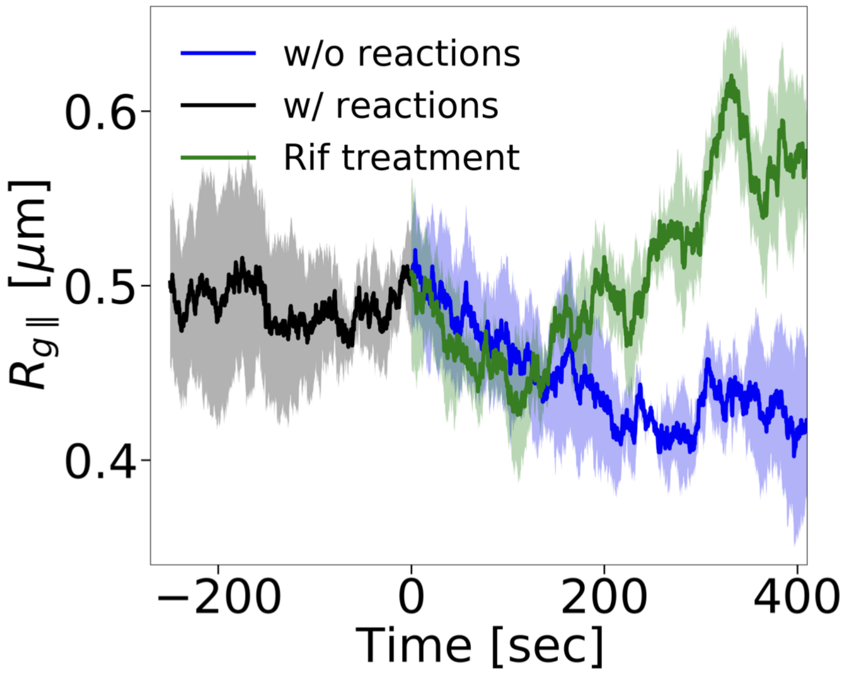


Figure S5: **Comparison of time-dependent nucleoid size, $R_{g\parallel}$, when all reactions are turned off versus during rifampicin treatment.** The blue curve represents the case where all reactions are disabled at $t = 0$, while the green curve represents the Rif treatment scenario, where only transcription and ribosomal treadmilling are inhibited. The black curve shows $R_{g\parallel}$ before reaction conditions are modified, that is when all reactions are present. The shaded area represents the standard error of the mean over three simulation runs.

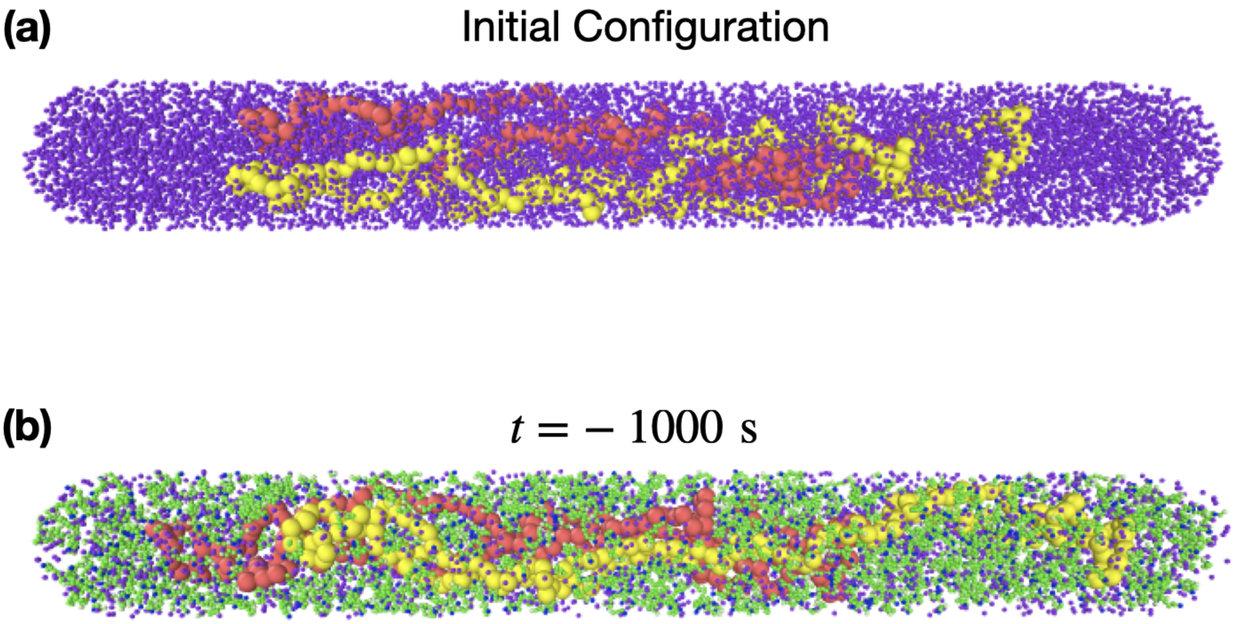


Figure S6: **Snapshots of the configurations in the two chromosomes simulations** (a) A snapshot of the initial configuration for the two chromosomes simulations. The initial state consists of two completely overlapping circular DNA polymers (red and yellow) surrounding with free ribosomes (purple). (b) A snapshot of two partially overlapping chromosomes at $t = -1000$ second (with respect to $t = 0$ as in Fig. 8(a) of the main text). Due to the out-of-equilibrium processes, most ofribosomes are part of polysomes (green and blue). For clarity, RNAPs are not shown.

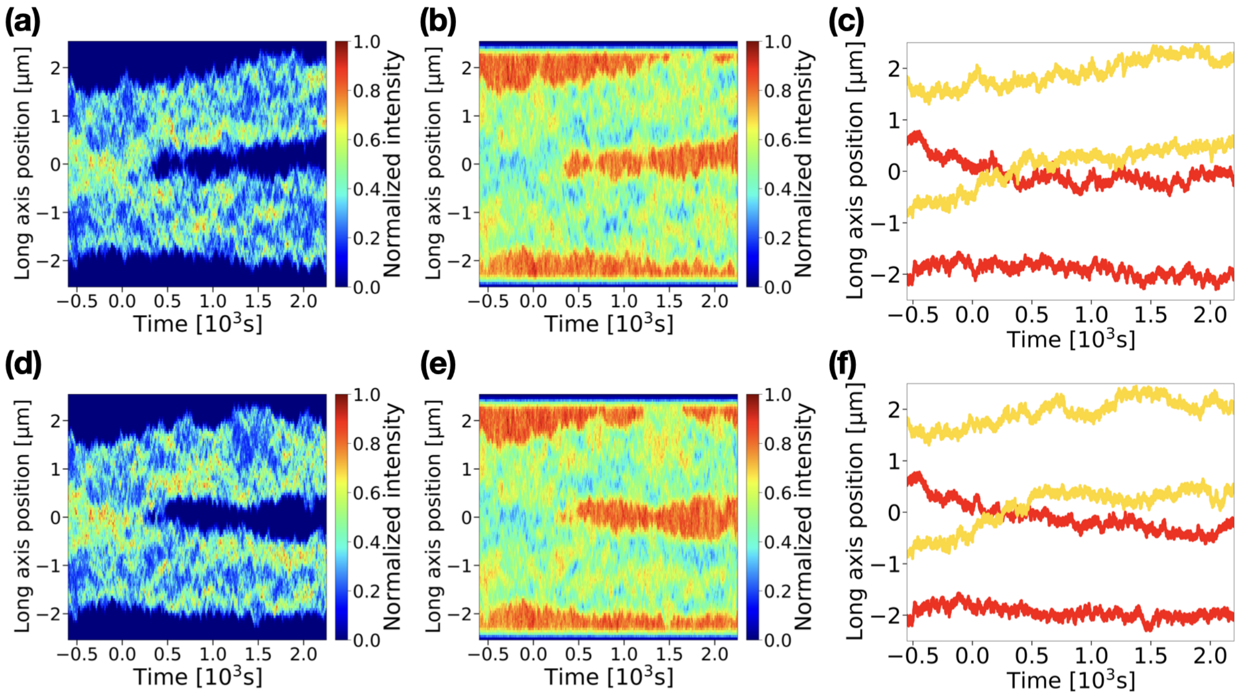


Figure S7: **Spatiotemporal distributions of two sister chromosomes and ribosomes under out-of-equilibrium conditions from two additional simulations that include out-of-equilibrium reactions** (a, d) Kymographs of the DNA density along the cell's long axis. (b, e) Kymographs of the ribosome density along the cell's long axis. (c, f) Axial positions of the outermost DNA monomers as a function of time. Red curves indicate the two outermost monomers of one DNA ($DNA_1$), and yellow curves another DNA molecule ($DNA_2$).

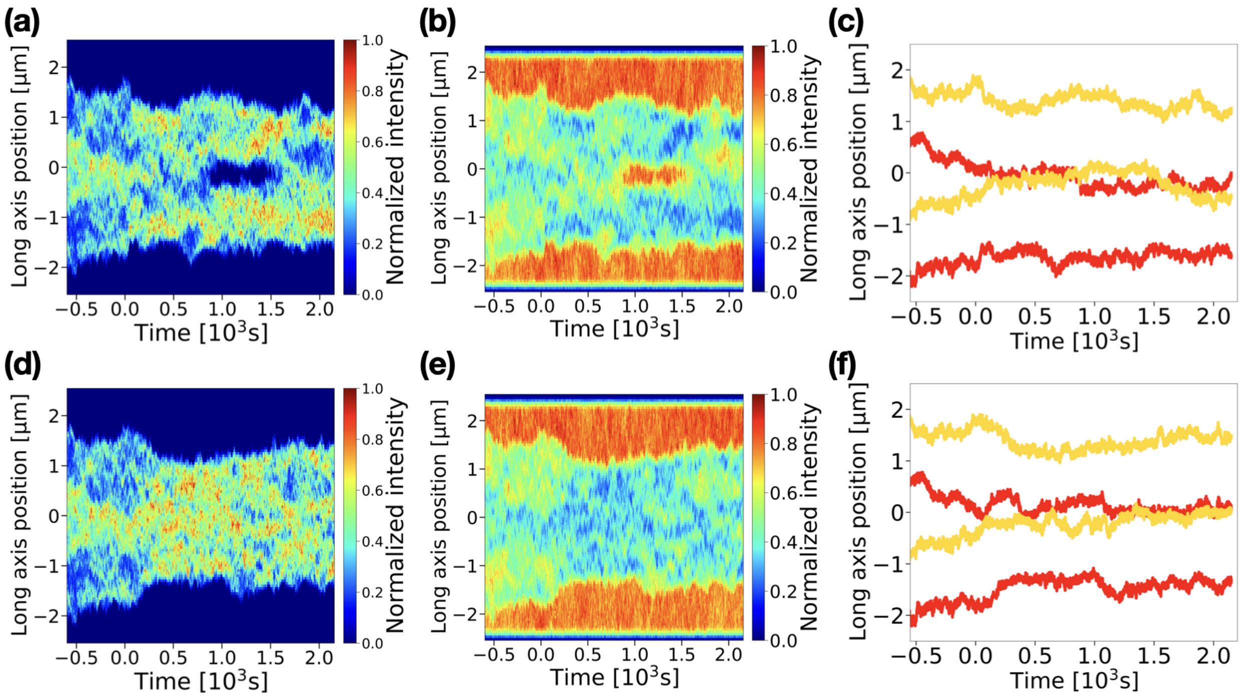


Figure S8: **Spatiotemporal distributions of two sister chromosomes and ribosomes under equilibrium conditions from two additional simulations** In these two simulations, all out-of-equilibrium processes are disabled at $t = 0$. (a, d) Kymographs of the DNA density along the cell's long axis. (b, e) Kymographs of the ribosome density along the cell's long axis. (c, f) Axial positions of the outermost DNA monomers as a function of time. Red curves indicate the two outermost monomers of one DNA ($\mathrm{DNA}_1$), and yellow curves another DNA molecule ($\mathrm{DNA}_2$).
